# Incorporating buccal mass planar mechanics and anatomical features improves neuromechanical modeling of *Aplysia* feeding behavior

**DOI:** 10.1101/2024.09.17.613591

**Authors:** Michael J. Bennington, Ashlee S. Liao, Ravesh Sukhnandan, Bidisha Kundu, Stephen M. Rogers, Jeffrey P. Gill, Jeffrey M. McManus, Gregory P. Sutton, Hillel J. Chiel, Victoria A. Webster-Wood

## Abstract

To understand how behaviors arise in animals, it is necessary to investigate both the neural circuits and the biomechanics of the periphery. A tractable model system for studying multifunctional control is the feeding apparatus of the marine mollusk *Aplysia californica*. Previous *in silico* and *in roboto* models have investigated how the nervous and muscular systems interact in this system. However, these models are still limited in their ability to match *in vivo* data both qualitatively and quantitatively. We introduce a new neuromechanical model of *Aplysia* feeding that combines a modified version of a previously developed neural model with a novel biomechanical model that better reflects the anatomy and kinematics of *Aplysia* feeding. The model was calibrated using a combination of previously measured biomechanical parameters and hand-tuning to behavioral data. Using this model, simulation feeding experiments were conducted, and the resulting behavioral metrics were compared to animal data. The model successfully produces three key behaviors seen in *Aplysia* and demonstrates a good quantitative agreement with biting and swallowing behaviors. Additional work is needed to match rejection behavior quantitatively and to reflect qualitative observations related to the relative contributions of two key muscles, the hinge and I3. Future improvements will focus on incorporating the effects of deformable 3D structures in the simulated buccal mass.

**Author summary:** Animals need to produce a wide array of behaviors so that they can adapt to changes in their environment. To understand how behaviors are performed, we need to understand how the brain and the body work together in their environment. One tractable system in which to study this brain-body relationship is the feeding behavior of the sea slug *Aplysia californica*. Despite having a small fraction of the number of neurons that humans have, this animal can produce many behaviors, respond to a changing environment, and learn from previous experiences. We have create an improved computer model of the slug’s mouthparts that simulates many of its key muscles and the forces they produce, together with a representation of the network of neurons that control them. With this model, we can recreate the feeding behaviors that we observe in the real animal, including biting, swallowing, and rejection, and use it to make quantitative predictions of how the animal will behave and respond to different stimuli. We found however that some aspects of the system were not well represented by simple 1-dimensional muscles, as has been done in most biomechanical models to date, but requires us to consider more complicated deformations of these soft bodies. Using this model as a tool, we aim to test hypotheses about brain-body interactions in the sea slug to better understand the behavior of small, slowly moving animals.

## Introduction

Animal behavior arises from the complex interactions between an animal’s nervous system, its muscles and their structural organization, and the environment [1, 2]. It remains an open question how these interactions produce behaviors; in addition, factors such as the size of the animal and the speed of movement profoundly shape the control challenges that brain and body must overcome [3, 4]. In addition to enforcing constraints, the body provides critical feedback that regulates and modulates circuits in the nervous system [5, 6]. Finally, the nervous system can exploit the architecture, compliance, and damping of biomechanical systems, which provide a morphological intelligence that reduces the required complexity of higher-level controllers [7, 8]. These factors all combine to suggest that the nervous system cannot be studied in isolation [1, 9], but rather in its appropriate mechanical and environmental context.

Brain-body interactions have been previously studied using neuromechanical models in many model organisms across phyla and in various behaviors. Walking and running are commonly studied, as they are vital to survival in most animals (e.g., fruit fly [10], cockroach [11], stick insect [12], rats [13], and humans [14]). Other modeling studies have examined grasping and manipulation [7], jumping [15], and swimming [16], amongst many other behaviors. In all of these systems, rigid skeletal components (e.g., bones and exoskeletons) provide the primary structure and constrain the system to a finite number of degrees of freedom. This allows modelers to take advantage of a host of well-established rigid body mechanics mathematical frameworks (e.g., Lagrangian mechanics, screw theory [17, 18], etc.) and computational tools (e.g., PyBullet [19], Mujoco [20], Animatlab [15], etc.). Additionally, the complex 3D geometries of muscles and other soft structures are often reduced to simple line elements, and their contact interactions neglected. This is often an acceptable simplification as the kinematics of these systems are predominantly dictated by the configuration of the rigid components and their articulations. As a consequence, the exact deformation of the muscles and soft elements may not be salient. These powerful modeling tools provide deep insights into the motor control of these endo- and exoskeletal systems. However, many animals lack significant rigid components. From nematodes and other worms to large cephalopods, animals across a large range of scales rely solely on soft tissues to locomote, feed, reproduce, and interact with the world, and even animals with rigid skeletons often contain soft-tissue motor systems (for example, the vertebrate tongue).

Studying movement in soft-bodied animals can be far more difficult than in animals with rigid skeletons and defined joints. Instead of the finite degrees of freedom associated with rigid body chains, soft-bodied systems have infinite degrees of freedom. A major challenge in modeling is to reduce this to a tractable dimensionality, through abstraction or simplifying assumptions. In addition, pronounced changes in the mechanical advantage and gross configuration of muscles caused by contact between multiple soft structural components must be dealt with. Not only are soft bodies a challenge to model, but often an even greater challenge to control. Despite these mechanical complexities, soft-bodied animals successfully perform numerous complex behaviors often with nervous systems comprised of relatively few neurons. How do the brain and body interact in these soft-bodied systems to create multifunctional behaviors in the face of their inherent mechanical complexity?

One tractable model with which to study the interplay between neural control and biomechanics in soft-bodied systems is the marine mollusk *Aplysia californica* (Fig. 1(a)) and its feeding organ, the buccal mass (Fig. 1(b)) [21–29]. *Aplysia* feeds on a variety of macroalgae (seaweed) [30], which it ingests as long strips of material, in contrast to the rasping from surfaces characteristic of most gastropods. The buccal mass is a fully soft system consisting of multiple interconnected muscular and cartilaginous structures [31]. The outer muscles of the buccal mass (Fig. 1(c-d)) form a tube-like structure that connects the lips of the animal to the esophagus. Within these outer muscles is a ball-like grasper, called the odontophore (Fig. 1(e)), whose muscles surround and articulate a toothed cartilaginous surface, called the radula (Fig. 1(f)). During feeding, the outer muscles of the buccal mass push the odontophore forward toward the lips (protraction) and backward toward the esophagus (retraction), while the muscles of the odontophore alternately open and close the radula on food [31]. Changing the phasing and relative magnitudes of these movements generates different feeding behaviors, the three best characterized of which are biting, swallowing, and rejection. Biting is an exploratory behavior, characterized by a strong protraction and weak retraction of the odontophore, where the animal attempts to grasp nearby food. Once the animal has successfully grasped food with its radula, it switches to a swallowing behavior, during which a strong retraction of the odontophore allows for food to be passed posteriorly to the esophagus. Finally, if the animal senses that what it has grasped is inedible, it can reverse the phasing of radular closing relative to biting and swallowing and instead push food out of the buccal mass in a rejection behavior [30, 32].

**Fig 1.**
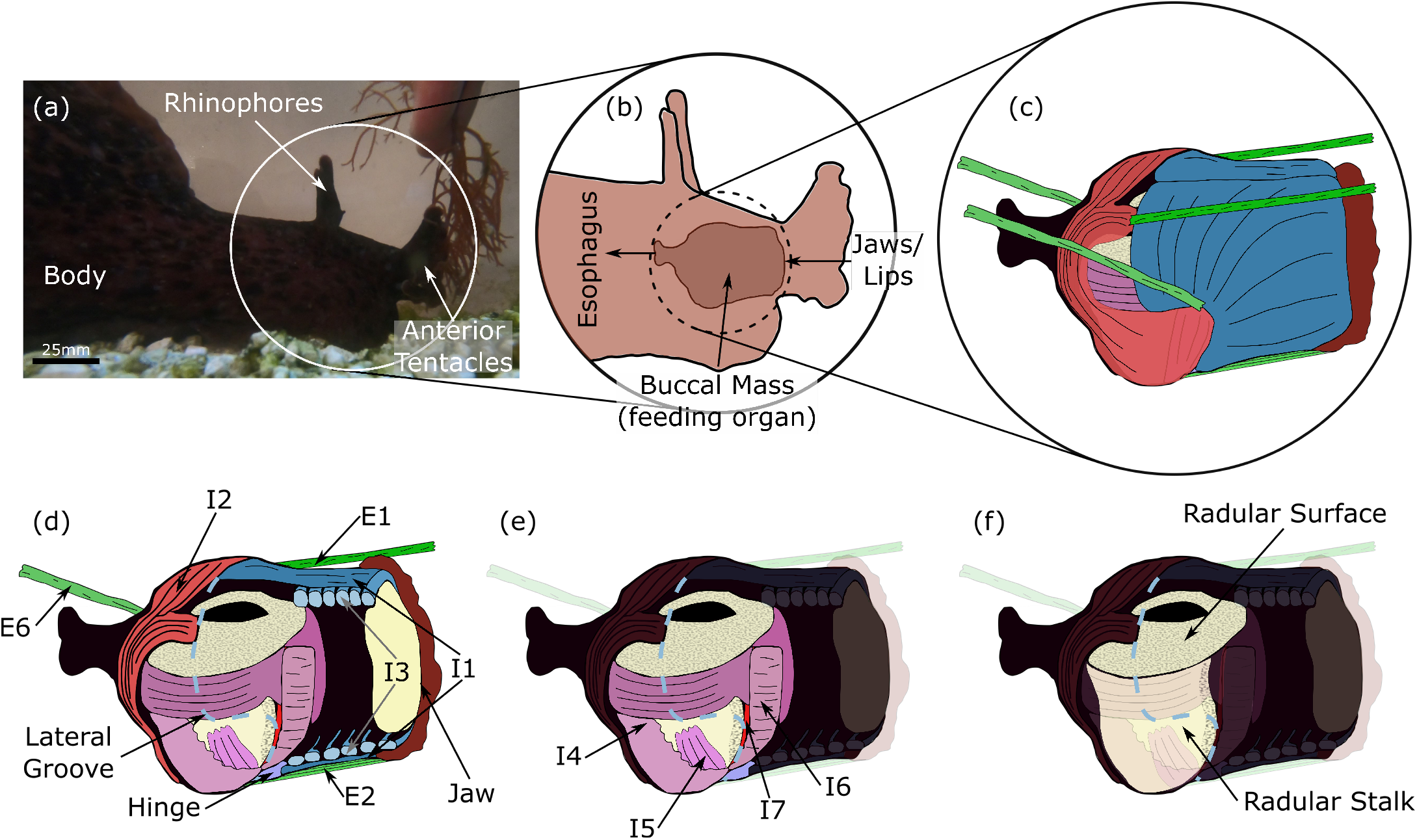
Overview of *Aplysia* feeding organ (buccal mass) anatomy. (a) An adult *Aplysia* feeding on *Gracilaria* seaweed. The white circle indicates the head of the animal, shown schematically in (b). The rhinophores and anterior tentacles both provide mechano- and chemosensory information to the animal. (b) Inside the head of the animal, the buccal mass connects anteriorly to the lips and posteriorly to the esophagus. Food enters through the lips and is carried through the buccal mass by the odontophore (grasper, (e)) where it is deposited into the esophagus. (c) False-colored anatomical drawing of the buccal mass musculature. All anatomical drawings ((c)-(f)) are modified with permission from [24]. The buccal mass is comprised of multiple interconnected muscles. (d) Cutaway anatomical drawing of the buccal mass showing internal structures. The outer muscles, which protract and retract the grasper, are shown labeled. Intrinsic muscles, which are wholly confined to the buccal mass, are designated “I” followed by a number and extrinsic muscles, which connect the buccal mass to the head, “E” followed by a number. (e) The internal musculature of the odontophore (grasper). (f) The grasping structure of the buccal mass is the radula (comprised of the radular stalk and radular surface), a tooth-covered cartilaginous structure articulated by the internal muscles of the odontophore.

The buccal mass neuromuscular system consists of a few key neurons that are large and identifiable that operate a limited number of muscles, which allows for behavior to be studied in a bottom-up fashion using *in vivo, in vitro, in silico*, and *in roboto* methods. Due to its slow behavioral cycling relative to its size, all feeding behaviors in the buccal mass remain quasi-static [22, 33], simplifying the analysis and modeling of the system’s mechanics since inertial forces can be neglected. This simplification applies across the entire size range of the animal [33], spanning from around 150 mg post-metamorphosis to over 1 kg in adults [34], meaning that the same model formulations can be used to model slugs of various sizes by making changes in model parameters.

Previous *in silico* and *in roboto* work has been dedicated to the modeling of *Aplysia*, at the component [35–37], subsystem [22, 23, 38], and system level [21, 24, 27, 39–42]. However, until now, models have either reduced the motion to single-axis translation [21–23, 40–42], which ignored behaviorally relevant kinematics [32], or were limited in their quantitative comparisons with *in vivo* data [24, 27, 39]. *In roboto* and other physical models are appealing for investigating the buccal mass because, being fully soft-bodied, the buccal mass can be difficult to model in its entirety. Robotic models allow for multibody physics to be captured in hardware, greatly reducing computational and mathematical complexity [43, 44]. However, *in roboto* models are limited by the available physical hardware and the accuracy of robotic components as analogs for the biological tissue. This is particularly true for soft robotic systems, which often must utilize bespoke actuator designs and fabrication modalities [45]. While previous *in roboto* models of *Aplysia* [24, 39] *have demonstrated the ability to produce multiple behaviors, there are still key discrepancies between their kinematics and those of the animal. In silico* models, on the other hand, can model individual components to arbitrary precision, restricted predominantly by the availability of experimental data for calibration and computational resources. However, with increased complexity comes rapid increases in computational and calibration costs, so a balance must be struck between computational speed and the model’s fidelity to the biological system.

In this work, we propose a new system-level neuromechanical model of the feeding system of *Aplysia*. This model incorporates additional biomechanical features that were previously neglected [21–24, 39] with the aim of reducing the error between modeled behavior and that observed in the animal while maintaining computational efficiency. A previously described Boolean nervous system model [21, 24] was modified to control a novel biomechanical model. This biomechanical model was developed using a demand-driven complexity framework, and thus, we added only the elements required to properly capture the animal’s behaviors. To this end, we hypothesized that the following simplifying assumptions (A) would still allow the model to adequately capture animal behaviors.

A1 Due to the bilateral symmetry of the system, the relevant mechanics of the system project fully to the midsagittal plane, and therefore, only 2D geometry and mechanics are required.
A2 The muscles and tissues of the system can be approximated using line element geometries.
A3 Bulk tissue passive forces do not play a significant role other than what can be captured through the above-mentioned line element structures and can thus be neglected in the model.

In the model presented here, additional anatomical features, including both muscles and passive elastic elements, were introduced to replace previously abstracted mechanical units. All model parameters are either implemented directly from previous experiments or hand-tuned to match existing kinematics data. The simulations produced by this model were then quantitatively compared with animal data, and areas of modeling mismatch were identified. With this model, we can better understand the mechanics of the buccal mass and how it generates adaptive feeding behaviors in *Aplysia*, and more generally, determine control principles applicable to other soft-bodied animals and how they may differ from animals with rigid skeletons. At the same time, the most significant deviations of the model from the biological system will determine what new components or degrees of freedom must be added to future models.

## Materials and methods

### Modeling

#### Relevant animal anatomy

Howells 1942 categorized the muscles of the buccal mass as either being intrinsic (I), entirely confined to the buccal mass, or extrinsic (E), connecting the buccal mass to the body wall. Muscles within the buccal mass can be roughly broken into two classes – the outer muscles that protract and retract the odontophore (Fig. 1(d)), and the muscles of the odontophore that articulate the radula (Fig. 1(e)) [31]. The anterior and posterior muscles interdigitate with each other and with the muscles of the odontophore at a region referred to as the lateral groove (shown as a light blue dashed line in Fig. 1(d-f)). The outer group of intrinsic muscles consists of the I1 and I3 muscles anteriorly and the I2 muscle posteriorly (Fig. 2(d)). The I1 is a thin muscle sheet that lies exterior and adheres tightly to the much larger underlying I3 muscle, which has muscle fiber orientations largely orthogonal to the I1. Together these muscles form a complex, with the I1 believed to be contributing to protraction [31] and the I3 providing the primary retraction force [46]. The I1/I3 is anchored anteriorly to the jaw (i.e. the most anterior region of the buccal mass that connects to the anterior body wall, Fig. 1(d)) [31]. The odontophore is moved through the lumen of the I1/I3 muscle complex during feeding behaviors.

**Fig 2.**
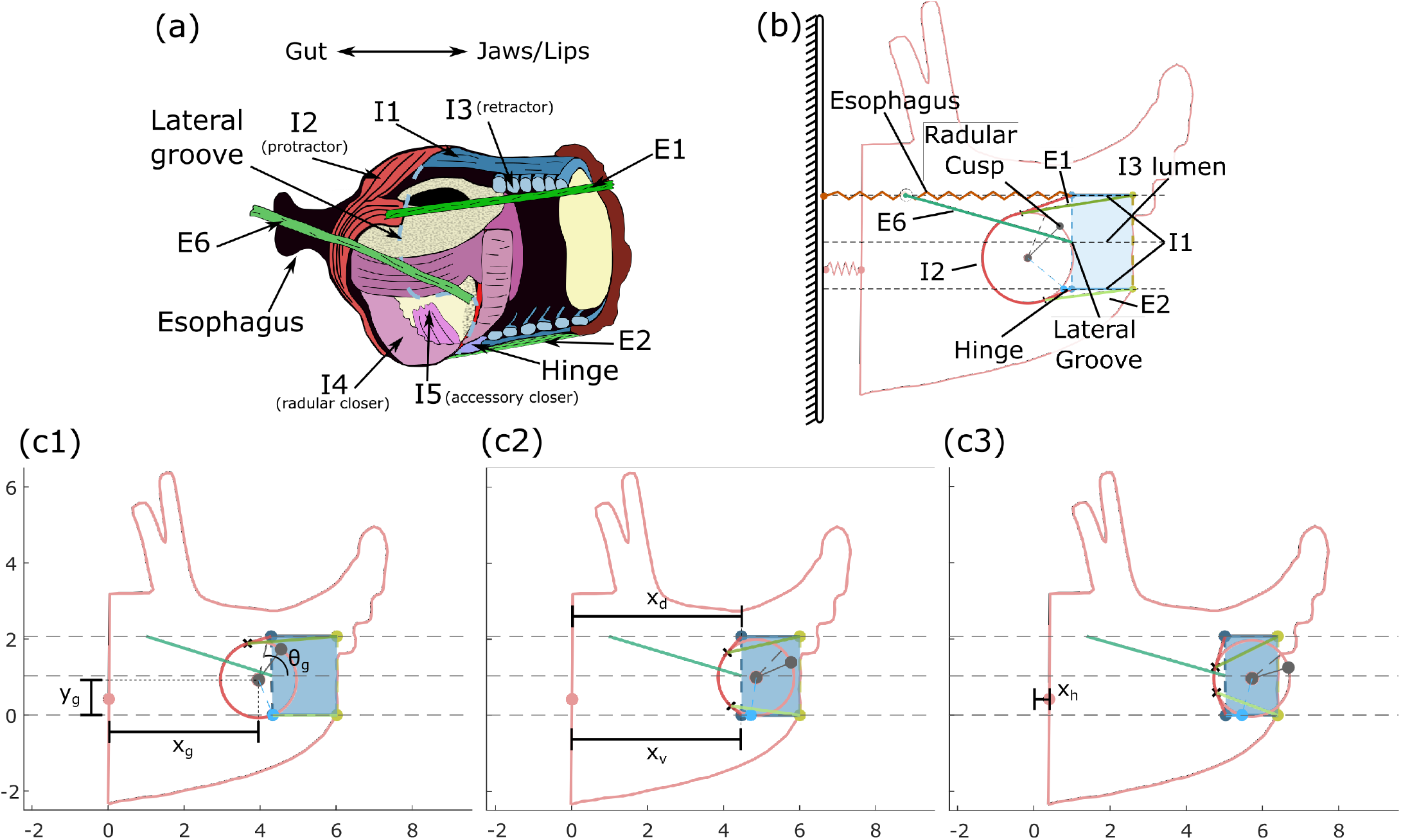
Comparison of (a) animal buccal musculature and (b) model anatomy. Analogous anatomical features share a color between (a) and (b). The outer intrinsic muscles (I1, I2, I3, and hinge) and the extrinsic muscles of the buccal mass are modelled using line element geometries. The odontophore is modelled as a rigid circle, with its musculature functionally abstracted as a single I4-like closer muscle. No grasper opener muscles were required for this model, and thus I7 was omitted. The rings of the I3 were modeled as an axial force acting at the I3 lumen midline but have no geometry to visualize. Note in (b) that the E6 attaches to the body wall at the same height as the esophagus, but the two structures do not interconnect. The point labelled as the “radular cusp” is where the model grabs food and represents the anterior-most point of the radular cleft (where food is grasped by the animal). In the model, we define the line from the radular cusp to the center of mass as the radular stalk axis. (c) Model degrees of freedom and multiple configurations. Frames (c1) - (c3) show three positions during the protraction of a biting behavior. Frame (c3) was slightly modified to ensure visibility of the *x*_*h*_ degree of freedom. Positions are in normalized model units. Panel (a) was modified with permission from [24].

The I2 muscle is another thin sheet that forms the back of the buccal mass, wrapping around the posterior of the odontophore and attaching both dorsally and ventrally to the I1/I3 complex at the lateral groove. The I2 serves as the primary protractor of the odontophore [29].

Several extrinsic muscles help to position the buccal mass inside the head [47]. These extrinsic muscles are anatomically (though not necessarily physiologically) more similar to muscles in rigid body systems, as they are mostly 1-dimensional elements connecting two points inside the animal. The E1, E2, and E6 extrinsic muscles can be represented as acting mostly in the midsagittal plane (Fig. 1(d)) and are thus relevant for this model. The E1 muscle connects posteriorly to the dorsal arms of the I2 muscle and anteriorly near the jaw. The E6 muscle attaches to the lateral groove and projects posteriorly to the body wall. The E2 muscle connects the ventral edge of the lateral groove to the jaw, running mostly parallel to the ventral I1 muscle. The E1 and E6 muscles contract during protraction, and E2 during retraction [48], though their contributions are not as critical to feeding effectiveness as the intrinsic muscles [47].

The odontophore muscles that articulate the radula consist of the I4-I10 muscles (Fig. 1(e), muscles I8-I10 not shown). The anatomy of these muscles and their functional roles in behavior are more complicated than the outer muscles that move the entire odontophore. In our model, the odontophore is assumed to be a rigid body (see “Model anatomy and degrees of freedom”), and thus the geometry of most of these muscles is not included. The functional role of the I4 muscles, which form much of the volume of the odontophore and act to close the radula on food [49], is included in the model, but no relevant geometry is included.

Finally, the hinge is an extended region of tissue formed by the interdigitation of the internal and external muscle, connecting the odontophore to the outer muscles of the buccal (Fig. 1(d)). The hinge connects the ventral edge of the lateral groove and has been hypothesized to assist in the retraction of the odontophore in biting and rejection [36].

#### Model anatomy and degrees of freedom

Key anatomical features of the musculature are included in our biomechanical model of the *Aplysia* buccal mass, which includes both intrinsic and extrinsic muscles (Fig. 2). From earlier models [21, 27], the odontophore is modeled as a rigid circle and is acted upon by forces generated by the I2 protractor muscle [29], the I3 retractor muscle [32, 50], and hinge muscle [36]. The I3 muscle is split into a bulk posterior portion that acts on the odontophore [50] and an anterior portion that generates a pinching force on food near the jaws [51]. The I4 muscle acts internally to the rigid odontophore to generate forces on strips of food [49]. In the model, the buccal mass sits inside a rigid head that is anchored to the global reference frame by a spring element representing the body wall of the animal.

To increase the anatomical accuracy of the model, the following features have been introduced: first, the I2 muscle was modeled as a chord that wraps conformally around the odontophore and attaches distally to a line (lateral groove) between the dorsal and ventral extents of the I3 lumen (Fig. 2b). Second, independent dorsal and ventral I1 muscles connect to the lateral groove and to the rigid head at the jaw. Third, in a previous model, the remaining muscles and connective tissue were previously represented as a single elastic element [21, 27]. To improve the biological fidelity of this model, this elastic element has been replaced by known muscles and tissues: a passive element representing the esophagus is attached between the dorsal extent of the lateral groove and the fixed reference frame; the E1, E2, and E6 extrinsic muscles were also added as active, contractile elements [47, 48]. E1 and E2 interdigitate with the I2 muscle and anchor at the jaw, whereas E6 attaches at the midpoint of the lateral groove and anchors posteriorly to the wall of the head. Additionally, the hinge muscle, previously modeled as acting at the odontophore center of mass [21], has been updated to better match the anatomy and has been modeled as attaching the ventrolateral odontophore to the ventral lateral groove. These anatomical features all capture some aspects of the true 3D geometry of the animal, but as part of our demand-driven complexity approach to this model, we hypothesize that the geometry of these muscular structures can be adequately captured using 1-dimensional line element geometries (A2).

To develop the equations of motion governing this system, the degrees of freedom (DoFs) must be defined. Though there are many distinct muscular and tissue elements in the system, many are geometrically constrained by one another. For this model, we further assume that due to the system’s bilateral symmetry, all relevant geometry and mechanics can be projected to the midsagittal plane of the head (A1). This removes two rotational DoFs and one translation DoF from each element. Finally, the head and the lateral groove are constrained to move only horizontally. Thus, six degrees of freedom remain (Fig. 2(c)) – the horizontal translation of the head (*x*_*h*_), the horizontal translation of the dorsal and ventral extents of the lateral groove (*x*_*d*_ and *x*_*v*_, respectively), and the horizontal and vertical center-of-mass translation and the rotation of the odontophore (*x*_*g*_, *y*_*g*_, and *θ*_*g*_, respectively). Here, *θ*_*g*_ defines the angle from the horizontal axis to the odontophore midline (Fig 2a). A scleronomic constraint is added to account for passive reactive forces from the I3 lumen and the hinge that are not explicitly modeled, following assumption A3. Specifically, the odontophore is constrained by a pin-slot joint, where a pin rigidly attached to the odontophore is free to translate in the horizontal direction and rotate in the slot but cannot translate in the vertical direction. This couples the center-of-mass vertical translation *y*_*g*_ and odontophore rotation *θ*_*g*_ by the kinematic constraint equation:

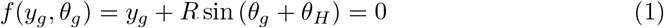

where *θ*_*H*_ is the fixed angle from the odontophore midline to the pin location on the odontophore, and *R* is the radius of the odontophore. In the animal, this coupling arises from complex interactions in a highly interdigitated structure. To fully model the interaction would require much more complicated mathematical methods that we deemed beyond the scope of our demand-driven complexity analysis. In addition to constraints imposed at the hinge, the system must be constrained to prevent interpenetration of the different muscular structures. Specifically, the odontophore must be prevented from passing through the ventral wall of the I3 lumen. In the animal, this arises naturally from these structures physically contacting each other; in the model, these constraints are captured by using rotational and translational inequality constraints (see “Penalty forces”).

#### Quasi-static equations of motion

Using the degrees of freedom defined above, the governing model equations can be derived using a Lagrangian framework, where gravity is omitted, and all passive elastic forces are included with active muscle forces in the sum of generalized forces. This reduces the Lagrangian to the kinetic energy of the system:

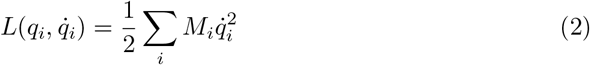

where *M*_*i*_ is the inertial parameter associated with the generalized coordinate *q*_*i*_. The generalized coordinates correspond with the degrees of freedom introduced above. Next, it was assumed that there exist non-specific damping elements in the system that can be captured by the dissipative potential:

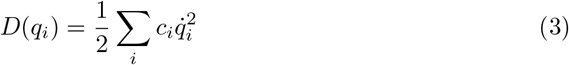

where *c*_*i*_ is the damping parameter associated with the *i*^*th*^ generalized coordinate. These damping parameters collectively abstract the viscoelasticity of tissues and any drag effects caused by the system being submerged in fluid. The units of *M*_*i*_ and *c*_*i*_ will depend on whether the generalized coordinate *i* is a length/spatial coordinate or an angle.

The influence of the pin-slot constraint, *f* (*y*_*g*_, *θ*_*g*_) (Eq 1), can be incorporated using a Lagrange multiplier *λ* representing a vertical reaction force on the odontophore pin. Because the additional rotational and translational constraints are inequality constraints and will therefore not be enforced throughout the cycle, here they are weakly enforced using penalty methods (See Penalty Forces below). Finally, the generalized forces associated with the *i*^*th*^ coordinate, accounting for both passive tissue and active muscle forces, are summarized as *Q*_*i*_. The model equations of motion can then be derived from the Euler-Lagrange equations as:

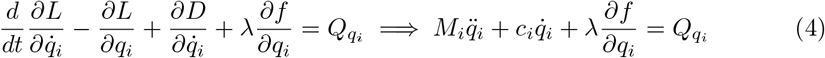

As previously observed, the inertia and accelerations in *Aplysia* feeding are several orders of magnitude smaller than other energetic sources and can thus be ignored [21, 22, 27, 52]. This yields the general form:

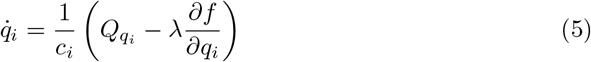

For *x*_*h*_, *x*_*v*_, *x*_*d*_, and *x*_*g*_, ∂*f/*∂*q*_*i*_ = 0, and the equation of motion (Eq 5) reduces to the scaled integration of the sum of forces. *y*_*g*_ and *θ*_*g*_ are influenced by the constraint equation (Eq 1), with *λ* specifically representing the vertical reaction force from the slot. The reaction force can be solved using the governing equation for *y*_*g*_, specifically:

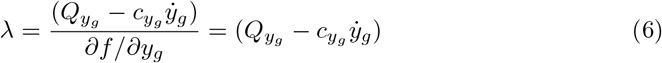

where 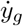 can be found by solving the constraint (Eq 1) for *y*_*g*_ and differentiating with respect to time:

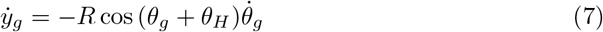

Substituting this and the derivative of the constraint (Eq 1) with respect to *θ*_*g*_:

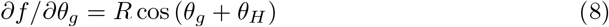

into Eq 5 for *q*_*i*_ = *θ*_*g*_ and solving for 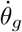 yields the governing equation:

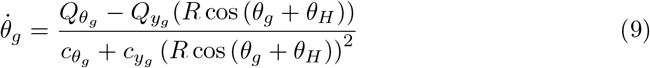

The governing equation for *y*_*g*_ then reduces to the derivative of the constraint equation (Eq 7). The components that sum together in *Q*_*i*_ are determined by the active and passive muscle forces of the model (Fig. 3f). These components are

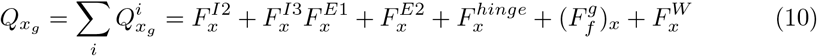

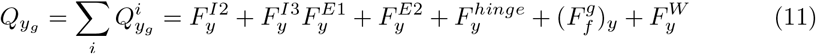

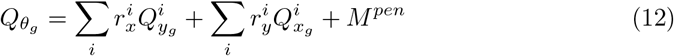

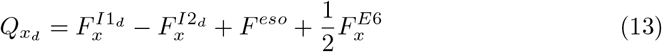

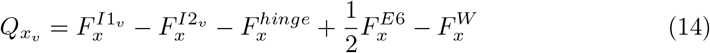

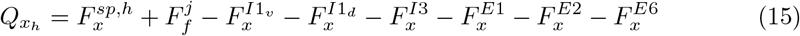

**Fig 3.**
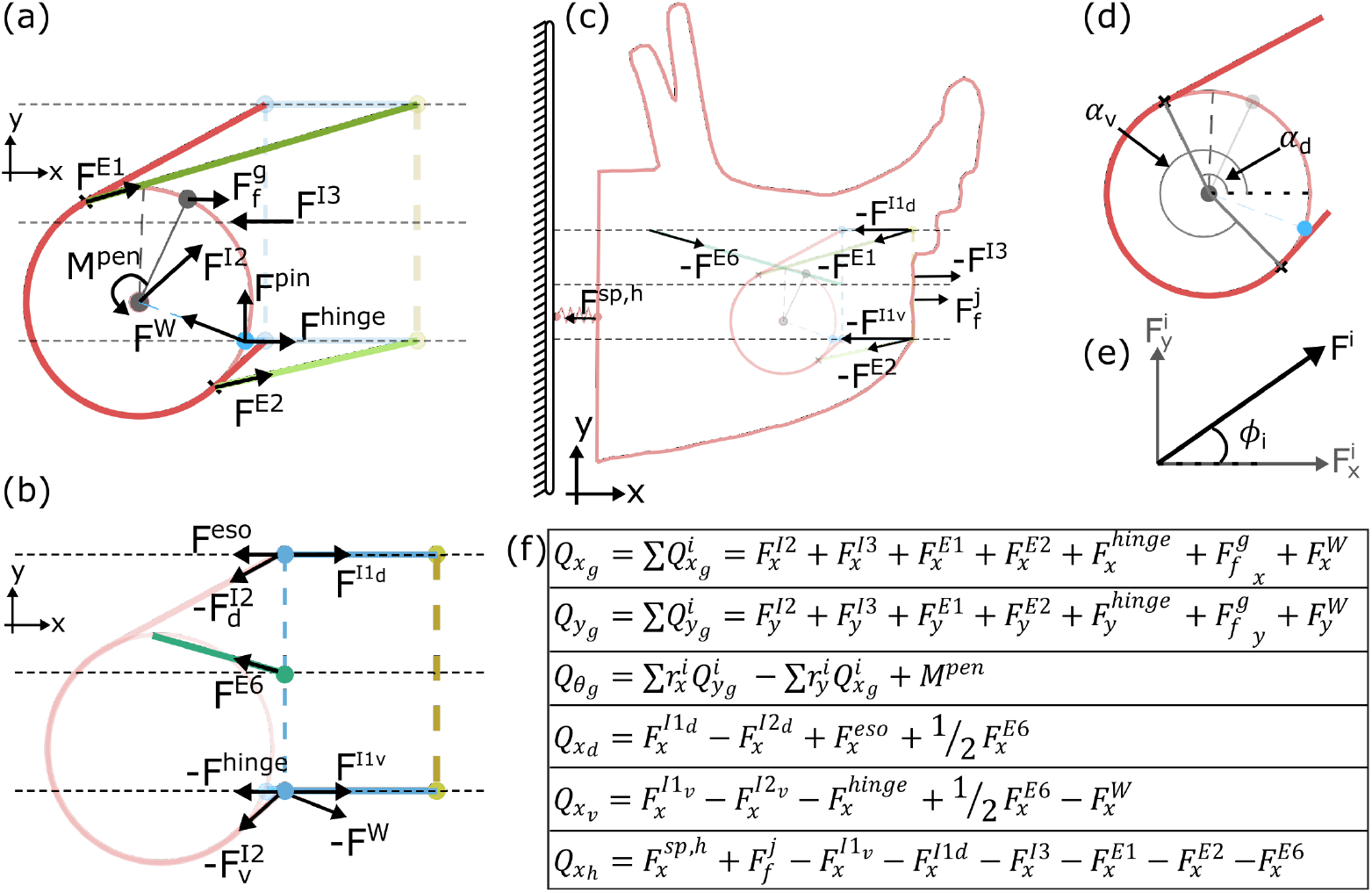
Summary of Model Mechanics. Forces exerted on the (a) odontophore, (b) dorsal and ventral lateral groove, and (c) head. Note, force arrows indicate the location of force application and *typical* direction of force application. All forces are defined as positive in the positive *x* and *y* directions, and the arrows drawn here do not indicate positive force direction. Arrow length also does not indicate force magnitude, as the magnitude will change throughout the feeding cycle. The force labeled *F* ^*pin*^ corresponds to the reaction force from the pin-slot joint equal to the Lagrange multiplier, *λ*. The olive dashed line indicates the jaw line (anterior edge of the I3 lumen) and where food is pinched by the anterior I3 muscle. (d) Definition of the tangency angles used to define the dorsal and ventral tangent points of the I2 muscle. (e) Definition of force decomposition and angle. (f) Summary of generalized forces associated with each coordinate. In the equation for the torques on the odontophore 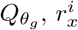 and 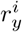 are the *x* and *y* components of the *i*^*th*^ force’s moment arm relative to the odontophore center of mass. In the equation for forces 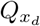 and 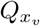, 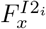, is the portion of the I2 force vector associated with the angle *α*_*i*_ (see Eqn. 23). The value *M* ^*pen*^ is the proportional penalty torque associated with the inequality rotation constraint, and *F* ^*W*^ is the contact penalty force associated with the ventral wall of the I3 lumen.

#### Muscle and tissue tensions and forces

The generalized forces in the equations of motion consist of the active muscle and the passive tissue forces. All muscles and springs, except for the dorsal and ventral I1, generate tension with positive magnitudes (no load in compression), and the geometry of the system determines the 2D force vectors that result from these tensions. The passive elements parallel with the I1 muscles can generate reaction forces in compression, representing the bulk elasticity of the I1/I3 complex. For simplicity, this is wrapped into the I1 dorsal and ventral forces. Passive tension in the *j*^*th*^ muscle/tissue is modeled as piecewise linear elastic elements governed by the equation:

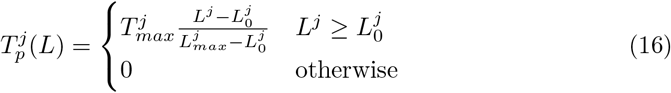

where 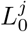 is the rest length of the *j*^*th*^ element and 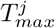(which has units of force) is the tension generated by the passive element at a length of 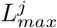 Here, *j* indicates the element’s identifier and is not an exponent. For the passive elements parallel to the I1 muscles, this conditional is dropped, and the tension is simply:

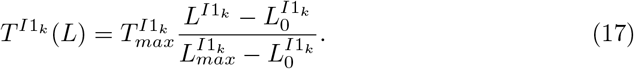

for *k*∈ [*v, d*]. The active forces generated by the muscles of the system come from a simplified muscle model taken from Webster-Wood *et al*. 2020 [21], where the active muscle tension *T* of the *j*^*th*^ muscle is the first-order filtered response of activation *A*, which in turn is the first-order filtered response of neural input *N*. This double-first-order filter is described by the following system of equations:

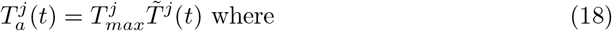

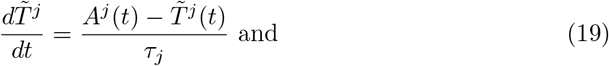

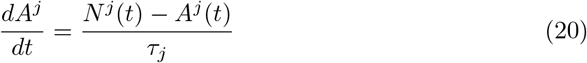

Here, *A*^*j*^(*t*) and *N* ^*j*^(*t*) are the normalized activation level *A*∈ [0, 1] and the neural input to the *j*^*th*^ muscle, and *τ*_*j*_ is the characteristic time constant associated with that muscle [21]. For all muscles except for I2, this is a fixed value. As in Webster-Wood *et al*. 2020 [21], the I2 has different time constants during activation and relaxation. The tension developed by the bulk posterior and anterior I3 muscle and the I4 muscle, comes only from this active component, as, in this model, there is no length associated with these muscles to generate passive forces. The tension in the I2, hinge, E1, E2, and E6 muscles are found as the sum of the active force model and the piecewise linear passive spring model:

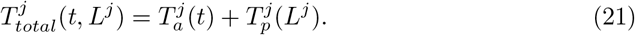

The tension in the I1 line elements is the same, save for the passive element being the fully linear model, as mentioned for Eq 17.

The resulting force vectors from these muscle and tissue tensions can be found using the model geometry and constraints. First, the anterior I3 and I4 muscle forces only pinch food and, therefore, do not need to be decomposed into a 2D force vector. Next, the dorsal/ventral I1 and hinge muscles are constrained to the *x* direction, so the *x* component of their force vector equals the tension in the muscle, and the *y* component is 0. The E1, E2, and E6 muscles act as simple line elements, and their forces can be decomposed as:

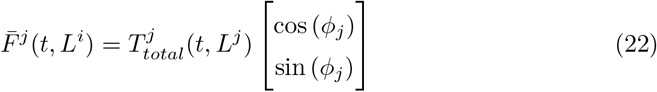

where *ϕ*_*j*_ is the angle that the *j*^*th*^ line element makes to the horizontal (Fig. 3e). This angle can be found knowing the fixed anchor and time-varying attachment points of the muscle. The attachment point for the E6 muscle is the lateral groove midpoint ([*x*_*d*_ + *x*_*v*_]*/*2, *H*_*lumen*_*/*2), where *H*_*lumen*_ is the height of the I1/I3 lumen. The attachment points for the E1 and E2 muscles are the points on the dorsal and ventral edges of the odontophore where the I2 becomes tangent, respectively. These points are also used to calculate the net force generated by the wrapped I2 muscle. At each point along the circumference where the I2 muscle is in contact with the odontophore, an infinitesimal force vector points inwards along the radius. Integrating these force vectors from the dorsal to the ventral tangency point yields the net force vector:

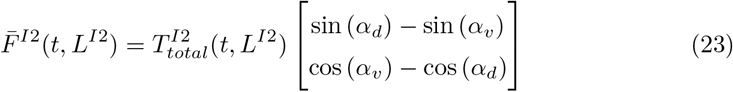

where *α*_*d*_ and *α*_*v*_ are the angles from the horizontal to the dorsal and ventral tangency points (Fig. 3d). These tangency points can be found from the odontophore center of mass location and the dorsal and ventral lateral groove points. The tangency point is defined by the point on the circle where the slope of the circle matches the slope of the line element running to the lateral groove anchor point. For the point *j* ∈ [*v, d*], let

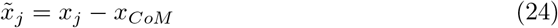

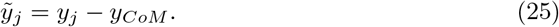

From these points, the tangency point 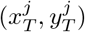 can be found as:

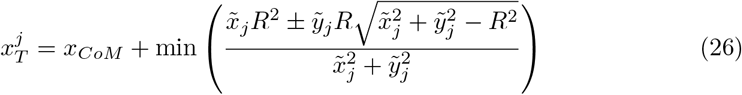

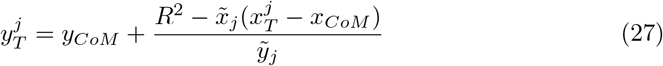

Using the min() function ensures that the solution to the quadratic problem corresponds to the point on the posterior edge of the odontophore.

Next, the hinge force vector only points along the x-axis, so the y-component of the vector is always 0. To reflect the observation that, at low stretches, the hinge is unable to generate force [36], the hinge tension is multiplied by a piecewise linear mechanical advantage function *MA*_*hinge*_. This function takes the form:

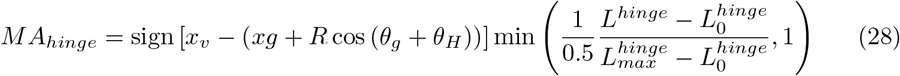

The sign() function term determines the direction of force application based on whether the odontophore hinge point is in front of or behind the lateral groove. The second term scales the tension linearly while the hinge length is less than 50% of its maximum length, after which point it scales the tension uniformly.

Finally, the force from the bulk I3 muscle is found as the tension in the bulk I3 muscle multiplied by a piecewise linear mechanical advantage function. This function reflects the fact that I3 applies force to the odontophore through a contact pressure and is thus related to the contact area between the odontophore and I3. This piecewise linear mechanical advantage (*MA*_*I*3_(*δ*)) equation yields a maximal force when the odontophore is fully internal to the I3 lumen and zero force when the odontophore is fully outside of the I3 lumen:

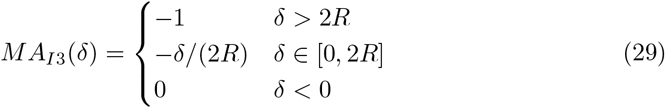

where

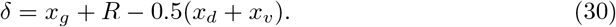

is the level of overlap between the odontophore and the I3 lumen. For the model presented here, the I3 force always acts along the horizontal lumen midline.

#### Penalty forces

As mentioned above, two inequality constraints (one translational and one rotational) were introduced to represent the contact forces coming from the I3 lumen which act to prevent the interpenetration of different structures. Because these constraints deal with inequalities (i.e., they are only enforced for particular configurations of the system), they cannot be handled directly using a Lagrange multiplier, as can the equality (pin-slot) constraint. Instead, these constraints are weakly enforced using linear penalty methods, where, if the governing inequality is not satisfied, a linear restoring force (or torque) is applied to the system in the direction that would act to enforce the constraint. For the rotational constraint, a torque is applied to the center of mass of the odontophore, but no reaction torque is applied to the I3 lumen as it has no rotational degree of freedom. The penalty torque is calculated as:

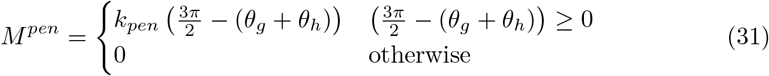

where *k*_*pen*_ is the stiffness of the penalty. The translational penalty force is applied if the point at *x*_*v*_ enters into the odontophore while the odontophore is outside of the I3 lumen. To check if the odontophore is inside the I3 lumen, we check if *x*_*g*_ *> x*_*hinge*_, which will only occur when the odontophore rotates into the I3 lumen. Here, *x*_*hinge*_ is the x position of the hinge point on the odontophore. To check if the point at *x*_*v*_ is inside the odontophore, we calculate the vector from the point at *x*_*v*_ to the center of mass of the odontophore. Let that vector be

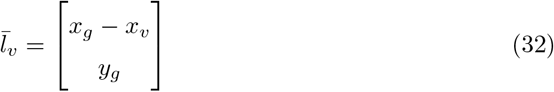

The force is applied if 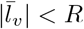. Therefore, the full penalty force is calculated as

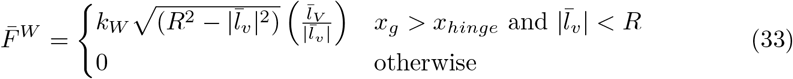

where *k*_*W*_ is the stiffness of the constraint.

### Frictional forces between the odontophore and seaweed

The model buccal mass can exert forces on seaweed using both its grasper and its jaws. In both cases, the force is applied through frictional contact. For simplicity, the velocity dependence of this frictional force [53] is ignored, and only static and Coulomb friction are considered, following the model presented in Webster-Wood *et al*. 2020 [21]. Briefly, for both the grasper (*g*) and the jaws (*j*), the frictional force (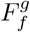 and 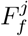respectively) is dependent on the state of slip. If the sum of other forces on the body in question is greater than the slip force, then the frictional force is determined by the grasping pressure and the coefficient of kinetic friction. The forces on the grasper and the jaws are, respectively, determined by the following:

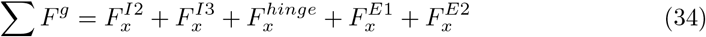

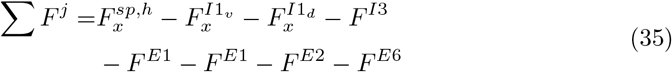

Note the force values 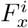 can be positive or negative depending on the current configuration of the grasper. The penalty force *F* ^*W*^ is omitted from this calculation as it only contributes during forward motion (when the force on food is 0). When the constraint is met, this force will be zero, and the inclusion of the force only impacts the calculation through the introduction of numerical noise. Most muscle forces acting on the head are subtracted because they are defined relative to the grasper and lateral groove and thus react negatively on the head. In addition to the forces on the body, the grasping pressure must also be determined individually for the grasper and the jaws. For the grasper, this grasping pressure *P* ^*g*^ is equal to *F* ^*I*4^, and for the jaws, the pressure *P* ^*j*^ is equal to 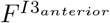 (the I4 muscle is internal to the odontophore (Fig. 2a) and is consequently not shown in Fig. 3).

If the sum of the forces is not greater than the slip force, then the force is equal and opposite to the other applied forces. For both grasper and jaws, this frictional force only acts in the horizontal direction, assuming that the anchor of the seaweed is infinitely far away. Because friction only acts in the horizontal direction, only forces in the horizontal direction are considered in the slip evaluation. Additionally, the seaweed is assumed to support no force in compression, so force from friction can never be negative. For the grasper (*g*) and the jaws (*j*), the slip condition becomes:

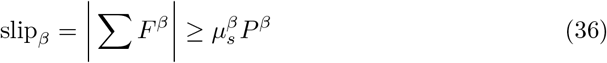

for *β* ∈ [*g, h*]. Together, with Eqs 34 and 35, the resulting friction force becomes:

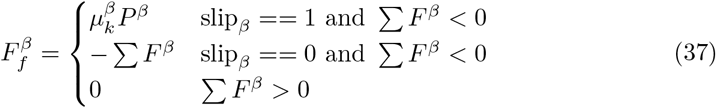

These forces are only applied if the seaweed is specified to be fixed (i.e. tethered and pulled taut). If the seaweed is specified to be detached, the frictional force is zero. For the case of no fixation, a secondary slip condition is tested for the grasper. Specifically, if 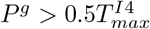, then the food is considered grasped (slip_*g*_ = 0). This indicator variable is used to calculate ingested/egested seaweed length (See “Model observables”). The jaw friction force is applied to the head, and the grasper friction force is applied to the grasper at a point defined by a fixed angle from the radular stalk axis, *θ*_*f*_. This point models the location of the cusp of the radular surface (Fig. 2) and is calculated as:

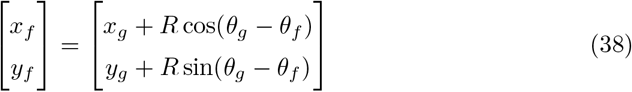

#### Neural controller

The neural controller utilized in this model is adapted from Webster-Wood *et al*. 2020 [21] and Dai *et al*. 2022 [24] (Fig. 4). Briefly, in the neural model, the activity of neurons is captured with Boolean variables, with 1 indicating spiking and 0 quiescence. The controller consists of three interconnected layers of neurons. Cerebral ganglion neurons integrate sensory cues to determine the behavior to be performed. Buccal ganglia interneurons then integrate cerebral neural and proprioceptive signals to determine the relative phasing of the model. Finally, buccal motor neurons integrate signals from the interneurons and cerebral neurons to activate the muscles in the biomechanical model.

**Fig 4.**
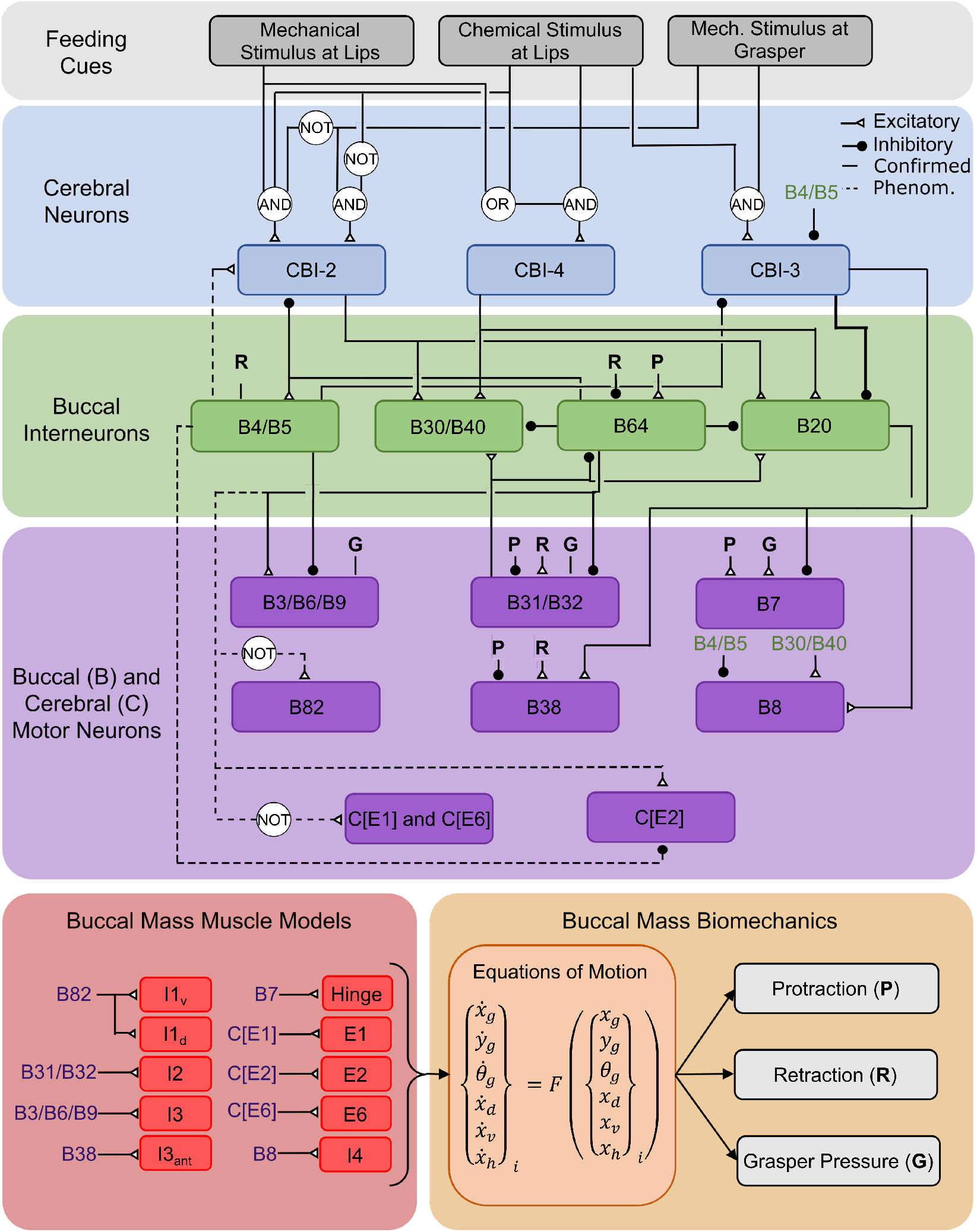
Neuromechanics Model Schematic. The neural circuitry of the buccal mass is modeled with a multilayered Boolean neural network modified from Webster-Wood *et al*. 2020 [21]. Here, each neuron is represented by a 1 or 0, with 1 indicating activity and 0 indicating quiescence. The user specifies feeding cues, and the cerebral buccal interneurons (CBIs) integrate these feeding cues to determine the behavior to be performed. The buccal interneurons then combine CBI and proprioceptive signals to determine the phasing of behaviors. Finally, buccal motor neurons combine interneuron and proprioceptive information to activate muscles in the biomechanical models. These muscles generate forces that influence the equations of motion (See “Quasi-static equations of motion”). Kinematic and kinetic variables are then calculated from this mechanical model to serve as proprioceptive signals. Dashed lines represent phenomenological connections within the network and may not represent true pathways. The bold line from CBI-3 represents an overriding inhibition that negates any other excitatory signals. As the motor neurons for the included extrinsic muscles have not been identified, but may reside in the cerebral ganglion with other extrinsic motor neurons [48], the motor neurons for extrinsic muscle Ex is labeled as C[Ex].

All neurons included in Webster-Wood *et al*. 2020 [21] are again included here, with additional neurons included for the muscles that were not present in previous models.

All additional connections introduced in this model are phenomenological and may not reflect true connections. In Dai *et al*. 2022 [24], the B43/B45 motor pool was included to activate the I1 muscles, with all I1 actuators sharing the same innervation. This motor pool was activated in retraction for all behaviors through connections to the CBI-3 cerebral neuron and the B64 interneuron. These connections were based on the assumption that I1 worked in tandem with the rest of the I1/I3 complex. However, closer examination of previous literature suggests that I1 and I3 have different functional roles. Church and Lloyd [50] described I1 motor neurons as causing “jaw shortening,” in contrast to I3 motor neurons causing “jaw closure.” Howells, in the original 1942 description of *Aplysia*’s feeding system [31], reported that I1 acts in conjunction with I2 and E1 to protract the odontophore. There is also evidence that the I1 motor neuron B45 may be doubly identified as B82 and that B82 can fire during the protraction phase [54]. Based on these findings, we hypothesize that I1 is activated during the protraction phase of the behavior and chose to model this protraction activity. The I1 motor neuron B43 is most active at the end of retraction after the cessation of the I3 retractor neurons [55]. Given that B43 activity comes after the major retractor motor neurons have stopped, it is possible that this activity also assists in protraction; i.e., helping return the odontophore to its rest position from its peak retraction. A complete description of the activity of the I1 motor pool requires further investigation beyond the scope of this work. Thus, in this model, we utilize B82 as the motor neuron for the dorsal and ventral I1 muscles and connect it such that it activates during protraction.

The extrinsic muscles E1, E2, and E6 were not present in either the previous *in silico* [21, 27] *or in roboto* [24] biomechanics models, and thus required additional model neurons. Previous animal studies identified E1 and E6 as protractor muscles [48], and thus, their motor neurons are active in protraction for all behaviors. E2 was identified as a retractor muscle [48] and is here activated in retraction for all behaviors. The motor neurons for these extrinsic muscles have not previously been identified in the literature, but the motor neurons of other extrinsic muscles have been shown to reside in the cerebral ganglion rather than the buccal ganglia [48]. Following the naming convention of other cerebral ganglion neurons, here, the motor neurons for the extrinsic muscle Ex are labeled as C[Ex].

The additional B44/B48 motor pool and the phenomenological motor pools RU1, RU2, and RU3, which separate out the components of the B3/B6/B9 motor pool introduced in Dai *et al*. 2022 [24] were not included here, as the muscles/muscle regions they innervate are not present in this model. Finally, as in previous model iterations [21, 24, 27], interneurons integrate proprioceptive feedback in the form of normalized grasper pressure and the relative position of the grasper in the head.Normalized grasper pressure 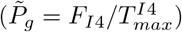 provides feedback about the closure state of the grasper. To allow for more direct quantitative comparisons with animal data [32], including quantifying translation in terms of the distance from the jaw line to the front of the odontophore, in this model, the relative odontophore position was calculated as:

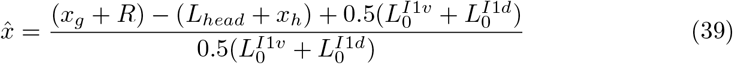

where *L*_*head*_ is the length of the rigid head, and *L*_*head*_ + *x*_*h*_ is the current position of the jaw line. The factor 0.5 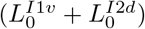 is the average length of the I3 lumen at rest and was incorporated to normalize this term to be 0 when the odontophore is fully retracted and 1 when the odontophore reaches the jaw line. *[ineq* is an affine transformation of the *x*_*gh*_ = *x*_*g*_ − *x*_*h*_ parameter used in Webster-Wood *et al*. 2020 [21] and is unitless.

### Computational methods and analysis

#### Model implementation

The model was implemented using Simulink (MATLAB r2022b, MathWorks). A variable step, variable order linear multistep numerical solver (*ode23*), was used to solve the dynamical system. Timesteps are restricted to be less than 10^−1^ s. For each timestep, the relative tolerance was set to 10^−5^, and the absolute tolerance was set to 10^−4^. To help minimize stiffness issues with friction models and abrupt changes of sign in muscle forces, zero-crossing detection was disabled. All simulations were run on an AMD Ryzen 5 5600X 6-core processor (3.70 GHz) with access to 16 GB of RAM.

#### Model observables

The state variables that are calculated while solving the model are not always directly comparable with animal data. However, numerical correlates of the measurements taken during animal experiments can be calculated from these states. For scalar model observables that deal with individual cycles, values were calculated only for steady-state cycles. For time-continuous observables, values can be calculated for any cycle. To obtain these steady-state cycles (which, for stable regions of the parameter space, occurred after one start-up cycle), simulations were run for 20 seconds, as this was sufficient for all behaviors to cycle multiple times. The beginnings of cycles were identified by the transition of the B31/B32 neuron from *off* to *on*. The model observables that were compared to animal data are summarized in Table 1. For details about how these observables were calculated, see Supporting Information 1.

**Table 1.**
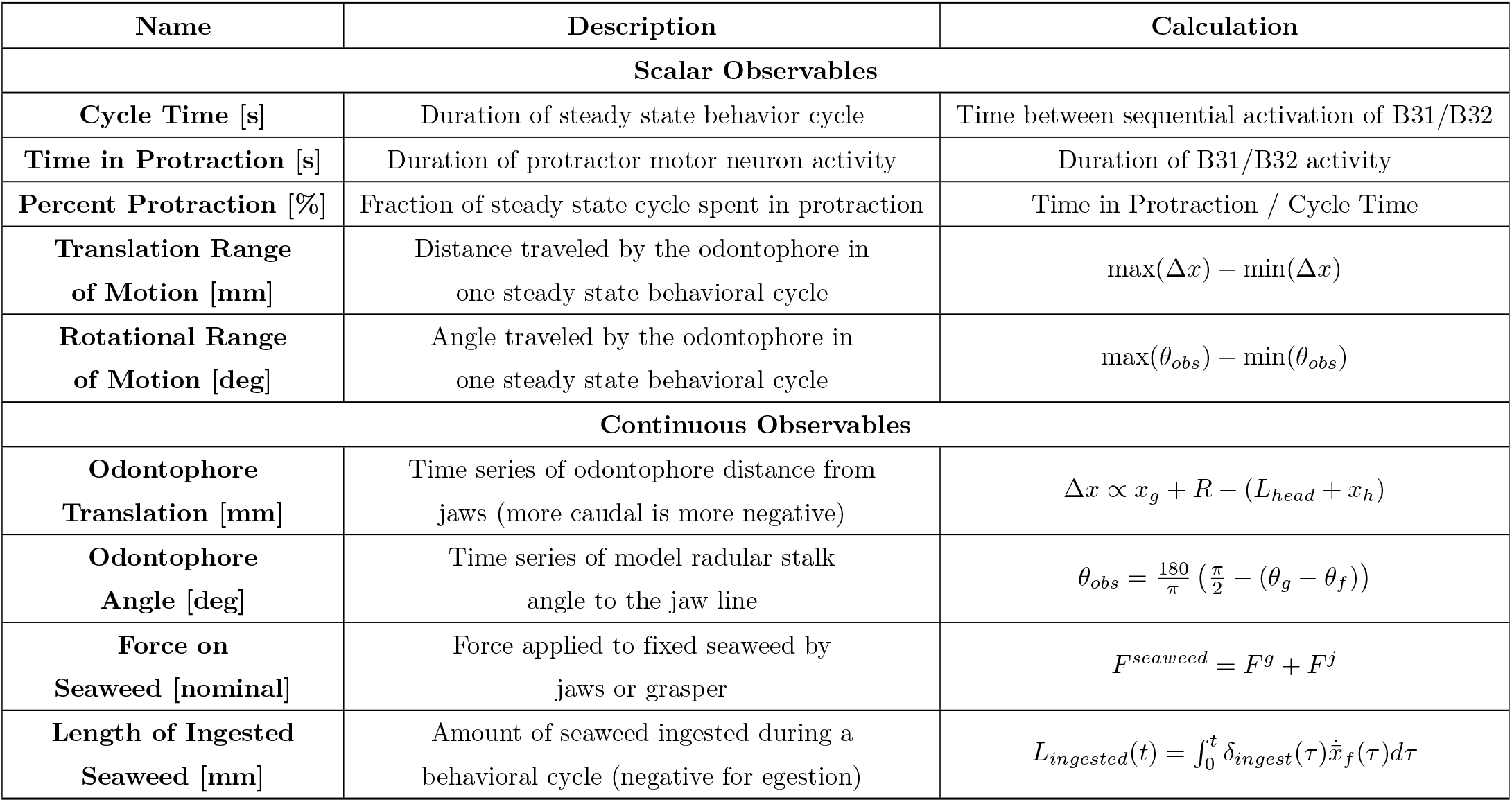
Summary of scalar and continuous model observables. For calculation details, see Supporting Information 1.

#### Parameter estimation and tuning

The model parameters were set using a combination of existing parameters from the literature, fitting submodels to animal data, and hand tuning. For the complete details of the model tuning, see Supporting Information 2. The anatomical parameters were calculated using the fixed buccal mass image presented in Neustadter *et al*. 2007 [32]. Muscle parameters for the I2, I1/I3, I4, and hinge were based on models and data from Yu *et al*. 1999 [35], Sukhnandan *et al*. 2024 [37], Morton *et al*. 1993, and Sutton *et al*. 2004, respectively. The remaining muscle parameters and the feedback thresholds in the neural model were hand tuned, first to produce multifunctional behaviors and then to match internal kinematic data. All timing parameters were normalized to the activation time constant of I2, and this value was set to achieve appropriate behavioral durations. All parameters used in this model can be found in the Supporting Information Section 4 and the provided code repository.

#### Animal data

Animal datasets were extracted and digitized from various literature sources to enable quantitative comparisons and assessment of model performance across all behaviors of interest. All time series data was digitized by hand using WebPlotDigitizer v4 [56]. Internal kinematics data were collected from Neustadter *et al*. 2007 for biting and swallowing [32] and from Novakovic *et al*. 2006 for rejection [23]. Neustadter *et al*. 2007 reports two different angles for the odontophore, one from the jaw line to the posterior edge of the I6 muscle, and one from the I6 to the radular stalk axis [32]. Because the model presented here does not have geometry corresponding to the I6, we cannot compare directly to these data. Therefore, we calculated the angle from the jaw line to the radular stalk axis from these animal data to compare to the model observables. These data correspond to cycles from 2 animals (total of 4 cycles). Novakovic *et al*. 2006 [23] reports a rotation for rejection, but which angle is reported is unclear. Therefore, only odontophore translation data were extracted from this paper. These data were from 1 cycle from 1 animal. To allow the kinematics data to be aligned and resampled for comparison with the model data, the animal data were fit to an 8^th^ order Fourier series. The Fourier series was fit to three concatenated duplicates of the single animal cycle to minimize the Gibbs phenomenon at the beginning and end of the cycles. This was performed for both the mean and standard deviation (if present) curves. These kinematic data from Neustadter *et al*. 2007 and Novakovic *et al*. 2006 were also used to calculate the translational and rotational range of motion (ROM) for the animal.

Neuromechanical data were extracted for swallows on untethered and tethered seaweed strips (referred to as unloaded and loaded swallows) from Gill *et al*. 2020 [26], and data for the length of ingested seaweed during swallowing was extracted from Lum *et al*. 2005 [28]. The neuromechanical data were generally not continuously periodic, with prolonged periods where the neural signals were zero, and the length of ingested seaweed data were strictly aperiodic. Therefore, a Fourier interpolation would not be appropriate. Instead, these data were spline interpolated to allow for resampling.

Timing and duration of protraction data for the different behaviors were collected from across multiple papers. Biting data was taken from Cullins et al. 2015 [57] (unpublished data) (*N* = 7 animals). Data for unloaded swallowing were taken from both Cullins *et al*. 2015 [57] and Gill *et al*. 2020 [26] (*N* = 12 animals total), and data for loaded swallowing were taken from Gill *et al*. 2020 [26] (*N* = 5 animals). For these datasets, mean and standard deviation data were available for multiple individual animals. To aggregate these data into single estimates of an “average *Aplysia*” in a way that incorporates the different number of samples per animal in the assembled datasets, a bootstrap resampling was conducted to obtain sampling distributions for the population-level mean and standard deviation (See Supporting Information 3.1). Data for rejection was digitized from plots of neural signals in Ye et al. 2006 [58] that aggregated data from 5 animals. Data were collected specifically for Type B rejections, as these are characterized by large egestion of seaweed similar to what is seen in the model. The start of the cycle was estimated as the beginning of I2 activity, and the end of the cycle was estimated to be the cessation of activity of the buccal nerve 2 (BN2) third largest unit [58], which likely reflects the activity of B43 [55]. The cycle length for rejection was estimated as the time between the start of I2 activity and the end of BN2 activity, with the standard deviation being calculated from the standard deviation in these values. The time at which I2 activity ended was also recorded to calculate the duration of protraction.

#### Statistical analysis

To validate this model with animal data and to investigate where additional model improvements may be needed, we conducted a quantitative comparison between the model observables and previously reported animal data. For time-varying signals, we calculated both the cross-correlation at zero lag and the root mean square error (RMSE). These signals include the translation and rotation kinematics for biting, unloaded swallowing, and rejection. We also calculated the cross-correlation between model and animal data for the swallowing neuromechanics reported by Gill *et al*. 2020 [26]. However, because the neural signals in the model and animal data report different quantities (neuron state for the model and median firing frequency for the animal data), the RMSE would not be meaningful. The cross-correlation was calculated at zero lag because signals were already aligned by either kinematically- or neuromechanically-relevant features in the signal. For the scalar observables, we were able to conduct statistical tests to compare model outcomes with animal data. When validating neuromechanical models, it is important for the predictions to be both statistically equivalent to and not statistically different from the animal data [59]. Both assessments are needed in this context because results being not statistically different does not inherently imply statistical equivalence [60]. Therefore, we conducted both t-tests to test for mean differences and equivalence tests, which we will now describe. One-sample tests were conducted because the model is fully deterministic and, therefore, has no distribution associated with its observables. For both test types, significance was set at *α* = 0.05. To take advantage of the bootstrapped distributions (See “Animal data”), we conducted the statistical tests based on the confidence interval on the metric of interest. For the t-test, the 95% confidence interval was calculated, and if the interval does not contain 0, then there exists evidence for a difference between the means at the 5% level [60]. For the equivalence test, the 90% confidence interval was calculated, and if the confidence was fully contained within a prespecified equivalence bound ([ − *d, d*]) expressed in the form of a Cohen’s d effect size, then this was taken as evidence that the means were equivalent up to a difference of *± d* [60]. We determined an appropriate equivalence bound by conducting a power analysis and choosing the bound that we could detect at a power of 1 − *β* = 0.8. For details of the power analysis, see Supporting Information 3.2. This analysis was conducted for each of the different behaviors because different sample sizes were available. For both loaded swallowing and rejection, there were data for *N* = 5 animals, so power was achieved at an equivalence bound of *d* = 1.65. For biting (*N* = 7), power was achieved at *d* = 1.27, and for unloaded swallowing (*N* = 12), power was achieved at *d* = 0.92. These were the bounds that were utilized for the equivalence tests in this work. These tests were conducted for Cycle Time and Time in Protraction, but not for Translational and Rotational Range of Motion, as these metrics only had data for either *N* = 1 or 2 animals.

## Code and data availability

All code and data are available for download on GitHub (Link to Model Code), and an archived version is available through Zenodo (doi: 10.5281/zenodo.13773332). All code is provided in Matlab r2022b.

## Results

### Model qualitatively produces multifunctional feeding behavior

Compared to previous neuromechanical models of *Aplysia* feeding, the biomechanical model presented here increases the degree of anatomical and mechanical agreement with the animal system. Compared to previous models, several previously abstracted structures have been explicitly modeled, and behaviorally relevant degrees of freedom, specifically the rotation of the odontophore and the length of the I3 lumen, have been incorporated. With this additional complexity, the model successfully reproduces the multifunctional behaviors demonstrated by *Aplysia* and by previous models when presented with different mechanical and chemical stimuli (Fig. 5). Biting responses were elicited when the model was presented with mechanical and chemical stimuli at the lips but not at the grasper. When the grasper was also presented with mechanical stimulus, the model instead performed a swallowing behavior. Finally, when the chemical stimulus was removed from the lips but a mechanical stimulus persisted in the grasper, instead of ingesting food, the model egested food in a rejection-like behavior. These behaviors were achieved by changing the chemical and mechanical sensory cues while only requiring a single, constant set of tuned parameters for the combined neuromuscular system.

**Fig 5.**
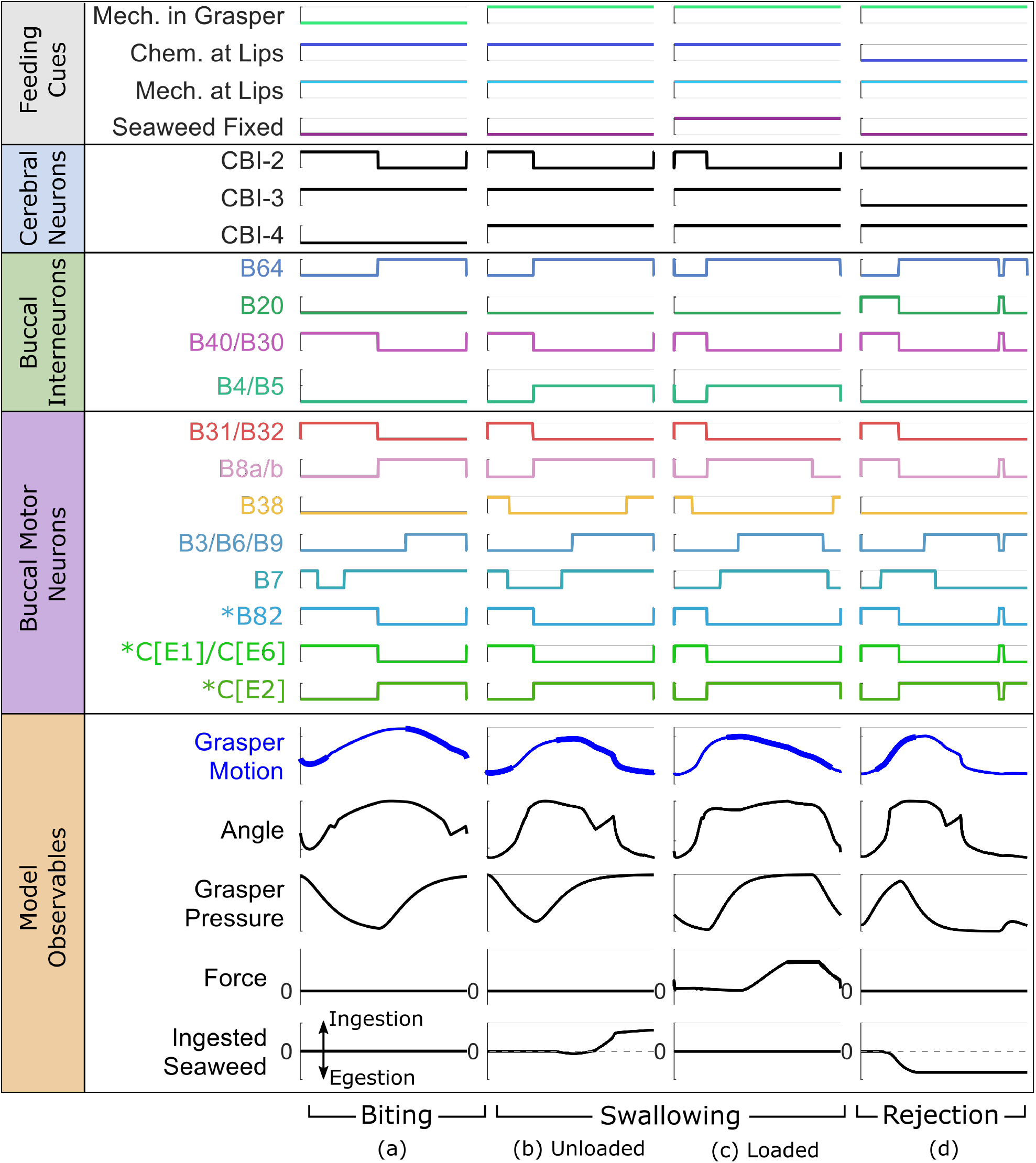
Steady state simulation results for (a) biting, (b) unloaded swallowing (untethered seaweed), (c) loaded swallowing (tethered seaweed pulled taut), and (d) rejection. Steady state was achieved after one start up cycle for all behaviors. Signals are grouped according to Fig. 4. Each behavior is generated by providing different feeding cues to the model. The bolded line in the Grasper Motion signal indicates when the grasper is closed 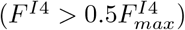. Dashed line in the Ingested Seaweed signal shows the level of no ingestion.

The model also successfully demonstrated the ability to adapt to changing environmental cues and maintain robust control of the feeding behaviors. This is seen in the model’s ability to change behaviors when sensory cues are switched mid-simulation (Fig. 6). During this simulation, the model is first presented with food but not allowed to grasp it (Phase A), during which time it performs a biting behavior. Then the model was allowed to begin pulling in the food (Phase B), at which point it switched to a swallowing behavior. As it continued to pull in food, the food was suddenly fixed in place (Phase C), and the model continued attempting to swallow, now generating force on the fixed food. Finally, the chemical cue indicating that the object on which it is feeding is edible was removed while a mechanical stimulus continued to be presented to the grasper (Phase D), at which point the model began rejecting the food and pushing it back out of the buccal mass. Behavioral robustness can also be seen in simulated feeding experiments where the food suddenly breaks and is no longer fixed (Fig. 7(a)). During these tests, the model begins feeding on unfixed seaweed. Then, at t = 2 s in the plots, the seaweed becomes fixed in place. However, unlike in previous simulations with fixed seaweed, here the seaweed has finite strength. If the force on the food exceeds that strength threshold, the food breaks and becomes unfixed again. Seaweed strengths were incrementally increased from breaking at 0 force to not breaking within the range of forces that the model generates on unbreakable seaweed. Throughout this range of strengths, the model could recover from the sudden change in boundary condition by the end of the current cycle and successfully switch back to an unloaded swallowing behavior. As with previous models [21], this is an emergent behavior of the network and not a programmed behavior. Additionally, as seen both in behaving animals [26] and in the previous model [61], with increasing strength of the seaweed comes an increase in the cycle length until it can no longer break the seaweed. However, the kinematics predicted by this model do differ during this break from the previous model. In the model presented in [21], the kinematics remain smooth through the transition from fixed food to unfixed food, with no sudden change in odontophore position. Here, the sudden change in force is accompanied by an initial sudden backward change in the odontophore position before returning to a smooth cycle. The magnitude of this change also increases with increasing seaweed strength. This difference is predominantly due to the increased damping in the previous model. Previously, the level of damping, characterized by the time constant of the dynamics, in normalized model units was set to 1. Here, the level of damping is much lower, set at a value of 1/50 in normalized model units. Thus we would expect the previous model to exhibit a far more damped response. Without access to internal kinematics data measured during similar types of experiments, we cannot ascertain whether this transition in the animal is rapid or smooth. If the transition in real animals is smooth, it may point to other mechanisms besides passive damping that act to decelerate the odontophore.

**Fig 6.**
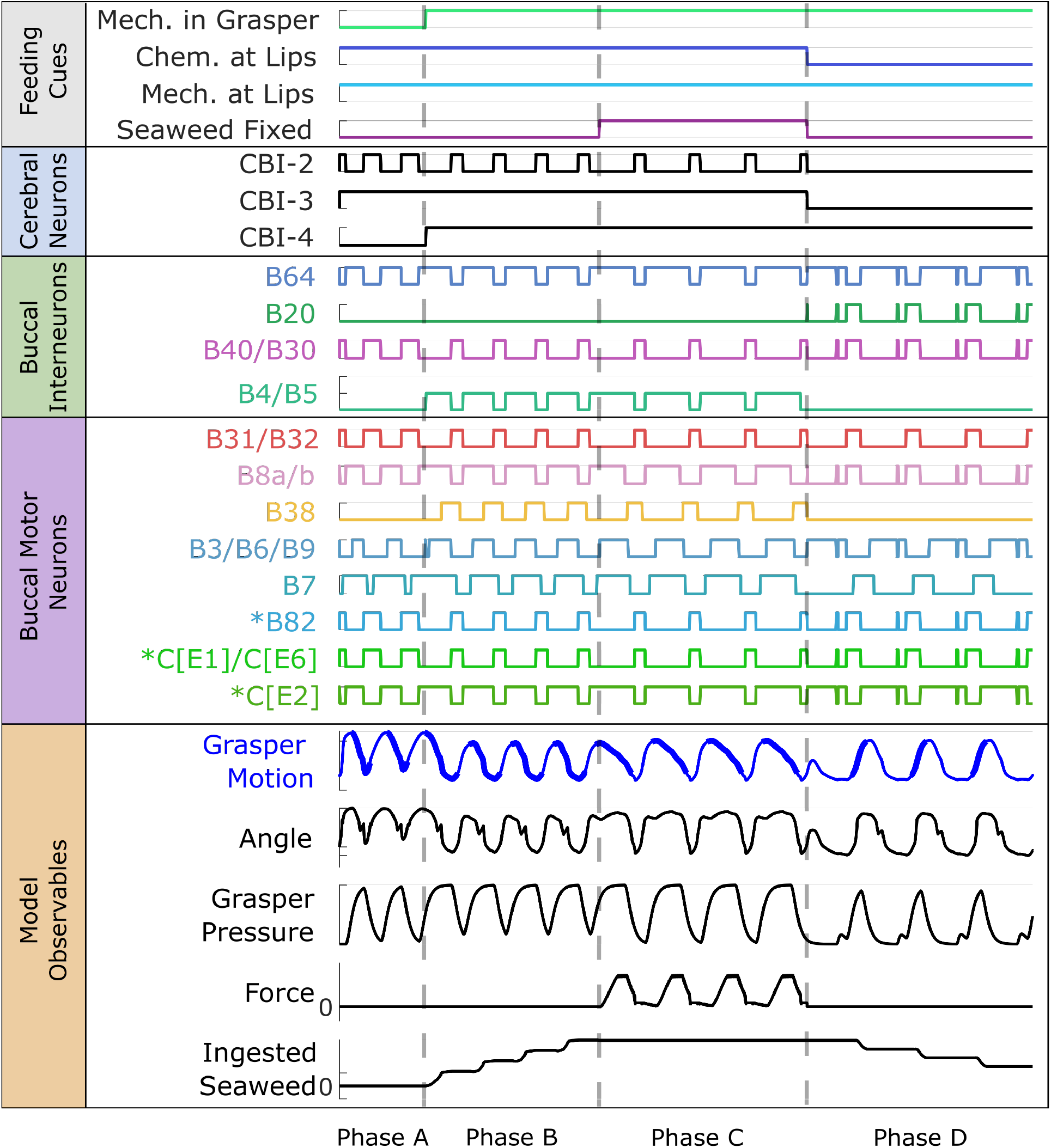
Behavioral switching. If the feeding cues are changed mid-simulation, the model can respond and successfully switch to a different steady-state behavior. For this simulation, the model is presented with food (Phase A), resulting in a biting behavior. Mechanical stimulus is then provided to the grasper (Phase B), causing a shift to an unloaded swallowing behavior. Then, the food becomes fixed (Phase C), and the model shows a loaded swallowing behavior. Finally, the chemical stimulus is removed from the lips of the model but mechanical stimuli continued to presented to the grasper (Phase D), and the model switches to a rejection behavior. Dashed gray lines show the dividing lines between the phases when the feeding cues are changed, but all results are from a single continuous simulation. Data is presented in the same manner as in Fig. 5 (see its caption for more details).

**Fig 7.**
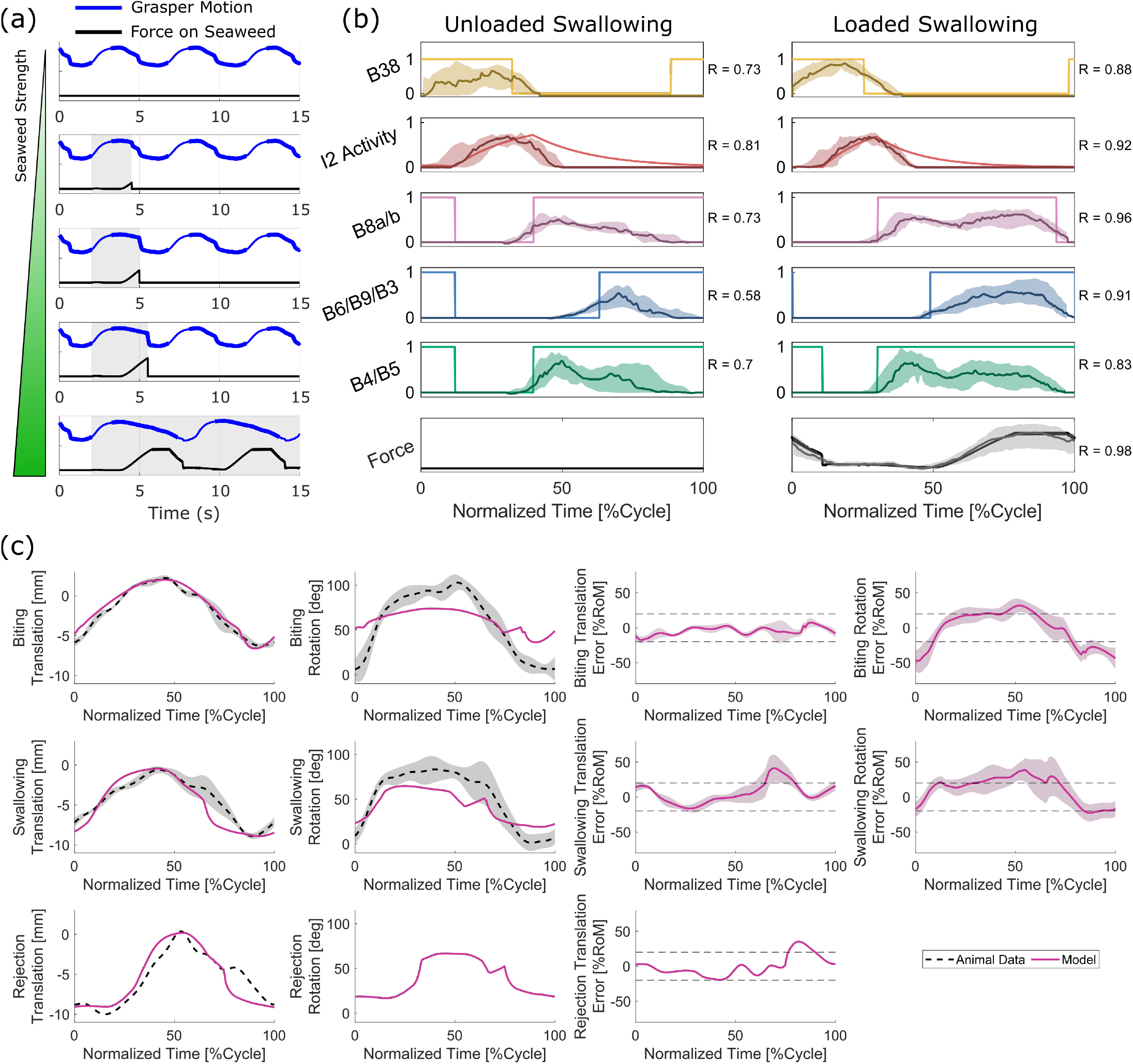
Computational experiments and comparison to animal data. (a) Feeding on finite strength seaweed. Each plot presents the grasper motion (blue) and force on seaweed (black) results for simulations with increasing strength of seaweed. Bolding of the grasper motion line indicates that the odontophore is closed, and the shaded gray region indicates when the seaweed is fixed (and not broken). As the force on seaweed reaches the failure strength of the seaweed, the food becomes unfixed and the model returns to an unloaded swallowing behavior. (b) Neuromechanics of unloaded and loaded swallowing behavior compared to animal data from Gill *et al*. 2020 [26]. Jagged lines and shaded regions show the median and interquartile range, respectively, of the animal data. The cross-correlation (R) was calculated for each signal. Signals were aligned to the beginning of I2 activity. (c) Comparison of model observables and internal kinematics as reported in Neustadter *et al. 2007* [32] for biting and swallowing and in Novakovic *et al*. 2006 [23] for rejection. Fuchsia lines show the model data, and black dashed lines and shaded regions show the mean *±* 1 standard deviation animal data. Columns 1 and 2 show the odontophore translation and rotation, respectively, for the model and animal. Columns 3 and 4 show the error between the model and animal kinematics, normalized to the range of motion (ROM), for translation and rotation, respectively. The gray dashed lines indicate an error of *±* 20% of the ROM. For rejection behaviors, only one translation cycle was available, so the standard deviation was not reported, and no rotation data was reported.

The model generally produces qualitatively similar steady-state results to those observed in animal experiments, providing support for the assumption that these behaviors can be captured solely through the midsagittal mechanics A1. However, there are two key areas where, even in preliminary comparison, we saw mismatches with the animal behavior, which contradict our assumption regarding the geometry of the buccal mass muscles (A2). First, during rejection behavior in the animal, it is seen that the multi-action neurons B4/B5 fire strongly at the onset of retraction [58]. This firing is stronger than their activity in other behaviors, which motivated the use of a ternary variable for its firing state in this and previous neural models [21, 24], instead of a standard binary state that is characteristic of a standard Boolean model. It has been demonstrated experimentally that this strong firing prevents the premature closing of the jaws and I3 muscle complex, which would prevent the successful retraction of the grasper in its open state [21, 58]. This retraction, therefore, would have to be mediated by a different muscle group, namely the hinge. However, B4/B5 does not fire during rejection behavior in the model using the current parameter tuning (Fig. 5(d)), and the brunt of retraction is still performed to the I3. If the tuning is changed so that B4/B5 fires during rejection, delaying the firing of B3/B6/B9, a much worse agreement was seen in the kinematics (Fig. 7(c)). The second mismatch also deals with an over-reliance on the I3 muscle group. During biting behaviors in animals, the motor neurons B6/B9 do fire during the retraction phase, but the frequency of firing is insufficient to generate significant forces in I3 [46]. Instead, retraction in biting may also be predominantly mediated again by the hinge [36]. In contrast, the model relies heavily on the I3 for generating retraction forces in biting, as seen by the long period of firing of B3/B6/B9 (Fig. 5(a)). If this firing is turned off in the model, the system does not cycle, and the odontophore becomes stuck in its forward position. These both point to shortcomings in the modeling of the hinge structure, which does not appear to be adequately approximated by simple line element geometries.

### Model comparison with animal data

The model was quantitatively and statistically compared to previously reported animal data using both time-varying and scalar metrics. The model agrees well with the internal kinematics reported in [32] (Table 2, Fig. 7(c)). For both translation and rotation in all behaviors, the cross-correlation was greater than 0.9, and the RMSE for translation was less than 1.5 mm. Additionally, for biting, the model was within *±* 20% of the range of motion (ROM) throughout the full feeding cycle (Fig. 7(c, Column 3)). This was also true for swallowing and rejection, except for a small portion of the cycle toward the end of retraction in both behaviors. In unloaded swallowing, 13% of the cycle had an error greater than 20%, with a maximum error of 41%. For rejection, 12% of the cycle exceeded 20% error, with a max error of 35%. However, the model dramatically underpredicts the rotational range of motion, particularly in biting, where the model achieved less than half of the rotational ROM of the animal. The agreement is better in swallowing but was still more than 2 standard deviations lower than observed in the animal. This comparison could not be conducted in rejection with the available animal data.

**Table 2.** Quantitative comparisons between model steady-state cycles and averaged animal kinematics [23, 32].

| Behavior | Cross-Correlation |  | Root Mean Squared Error |  |
| --- | --- | --- | --- | --- |
|  | Translation | Rotation | Translation [mm] | Rotation [deg] |
| Biting | 0.985 | 0.937 | 0.609 | 24.1 |
| Unloaded Swallowing | 0.979 | 0.978 | 1.31 | 18.1 |
| Rejection | 0.974 | — | 1.48 | — |
|  | Translational Range of Motion [mm] |  | Rotational Range of Motion [deg] |  |
|  | Model | Animal Mean [STD] | Model | Animal Mean [STD] |
| Biting | 8.61 | 8.50 [0.45] | 37.7 | 96 [15] |
| Unloaded Swallowing | 8.45 | 8.29 [0.57] | 45.7 | 81 [17] |
| Rejection | 9.30 | 10.4 [ ] | — | — |

For unloaded and loaded swallowing, the neuromechanics of the model were compared with data from Gill *et al*. 2020 [26], and the cross-correlation was calculated (Fig. 7(b)). In general, the cross-correlations were high, with all but one of the signals scoring *R* ≥ 0.7. The largest disagreements exist in the B6/B9/B3 signal in unloaded swallowing (*R* = 0.58). This discrepancy relates to the model being under-constrained in protraction. To achieve the correct kinematics, the parameters had to be tuned such that the B31/B32 motor neurons, which innervate I2, turned off partway through protraction. This allowed the odontophore to slow down before reaching peak protraction, at which point the B6/B9/B3 motor neurons activated generating force in I3 for retraction. Otherwise, the slope of the translation would be too high. However, for this to occur and for the odontophore to still achieve the appropriate range of motion, the firing of the I3 motor neurons needed to be delayed. If there were other mechanisms to slow down protraction, this delay would not need to occur.

The model was also compared to external data for swallowing behavior in the form of the length of seaweed that was ingested in a single swallow [28] (Fig. 8). Here, the model performed very well, with a cross-correlation of *R* = 0.996 and an RMSE of 0.746 mm. The model was not explicitly tuned to match this curve, but rather, this agreement arose because of the timing and kinematic agreement with animal data. As in the animal, the position of the seaweed does not change during the beginning of protraction, as it is held in place by the jaws (i.e., the pinched anterior of the I3 muscle). Near the peak of protraction, the jaws open, and food is slightly egested before the odontophore changes direction. At this point in the animal data, the food is pulled in towards the esophagus at a roughly constant rate before coming to rest at peak retraction. The rate of seaweed ingestion in the model is not as uniform as in the animal, with a period of more rapid ingestion followed by a period of very slow ingestion before coming to rest. Despite these differences, the model approximates the timing of seaweed ingestion and the length of seaweed ingested within a single swallow well.

**Fig 8.**
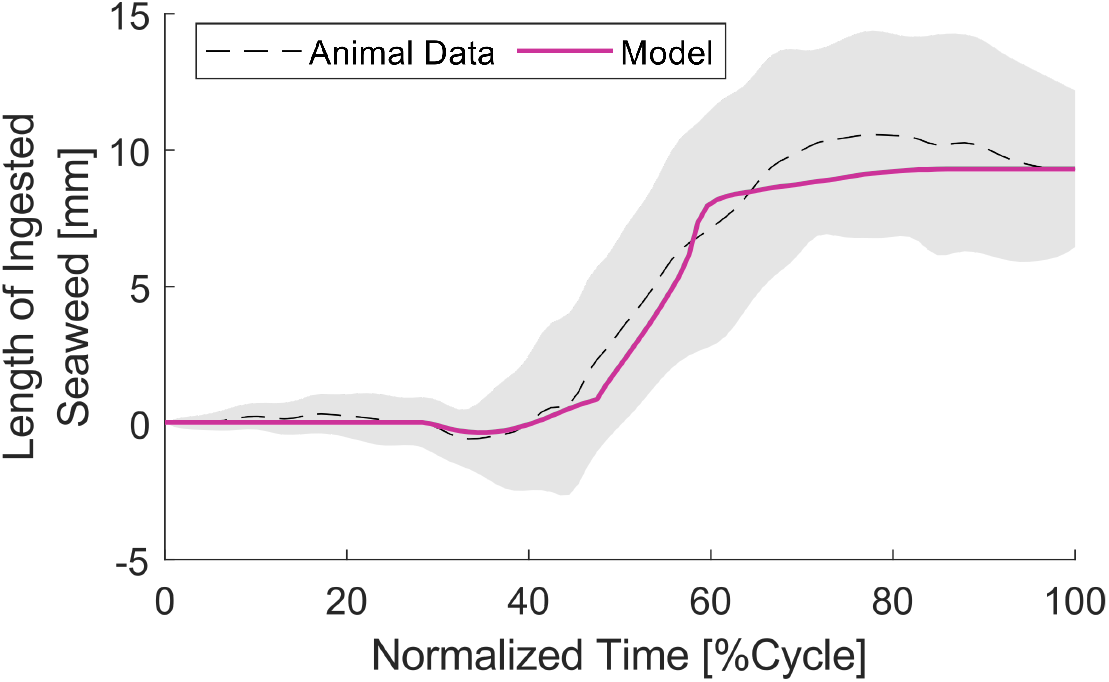
Length of seaweed ingested in a single swallowing behavior. The gray dashed line and shaded region show the animal data extracted from Lum *et al*. 2005 [28], and the fuchsia line shows the model data.

Finally, for all behaviors of interest, a statistical comparison was conducted for the length of behaviors and the fraction of behavior that was spent in protraction (Table 3, Fig. 9. Specifically, both equivalence tests and t-tests were conducted to determine if the model predictions agree with the animal data. For loaded swallowing, the raw cycle time was not compared to animal data, as the animals in this dataset skewed towards longer unloaded swallows and thus may not be representative of the whole population (See Supporting Information 3.3). However, the length of loaded swallows relative to unloaded swallows for a given animal may be a more consistent metric, so this was compared to model predictions. For all metrics related to biting, unloaded swallowing, and loaded swallowing, the model was statistically equivalent to and not statistically different from the animal data (Fig. 9, Table 3), suggesting that this model adequately represents an “average *Aplysia*” in these behaviors. However, both metrics for rejection were non-equivalent, with both the cycle duration and percent protraction being significantly lower than the mean animal value. This disagreement relates to the underperformance of the hinge and the under-constraining of the odontophore in protraction. As with swallowing behaviors, the rate of protraction was too fast compared to animal data. To achieve the correct time-normalized kinematics, the rate of retraction had to be similarly increased. This was achieved by activating the I3 muscle at the peak of protraction rather than waiting for the hinge to initiate retraction, as is seen in the animal [58]. If B4/B5 were to fire, delaying the firing of the B3/B6/B9 motor pool, the odontophore would sit near peak protraction for a prolonged period, only moving slightly because of how little force the hinge could generate. This would allow the cycle duration of rejection to better approximate the measured value but at the cost of a much worse kinematic match. The under-constraining of protraction in this model provides evidence against our assumption that the bulk tissue passive forces may be neglected (A3).

**Table 3.** Scalar metric comparison with animal data. Animal data were aggregated from previous literature [26, 57, 58]. Positive percent differences indicate that the animal mean is greater than the model value. The Equivalence column presents the results of the equivalence test (Y: 90% confidence interval is fully within the equivalence bound, see Fig. 9), and Difference column presents the results of the t-test (Y: 95% confidence interval does not intersect a standard difference of 0).

| Behavior | Animal Value<br>(Mean ± SEM)<br>[STD] | Number of<br>Animals | Model Value | %Difference<br>[Effect Size] | Equivalence | Difference |
| --- | --- | --- | --- | --- | --- | --- |
| Cycle Time [s] |  |  |  |  |  |  |
| Biting | 4.36±0.30<br>[0.77] | 7 | 4.38 | -0.36<br>[-0.02] | Y | N |
| Unloaded<br>Swallowing | 4.84±0.29<br>[0.98] | 12 | 4.87 | -0.54<br>[-0.03] | Y | N |
| Loaded<br>Swallowing | — | — | 6.37 | — | — | — |
| Rejection | 8.05±0.38<br>[0.85] | 5 | 6.99 | 13.2<br>[1.24] | N | Y |
| Percent Protraction [%] |  |  |  |  |  |  |
| Biting | 48.3±2.4<br>[5.9] | 7 | 46.4 | 3.92<br>[0.32] | Y | N |
| Unloaded<br>Swallowing | 28.7±2.4<br>[7.6] | 12 | 27.4 | 4.32<br>[0.16] | Y | N |
| Loaded<br>Swallowing | 18.5±1.8<br>[3.8] | 5 | 18.9 | -2.07<br>[-0.10] | Y | N |
| Rejection | 37.3±3.4<br>[7.6] | 5 | 23.6 | 36.8<br>[1.80] | N | Y |
| %Increase in Cycle Length (relative to unloaded swallowing) |  |  |  |  |  |  |
| Loaded<br>Swallowing | 32.9±10.1<br>[20.9] | 5 | 30.8 | 6.31<br>[0.10] | Y | N |

**Fig 9.**
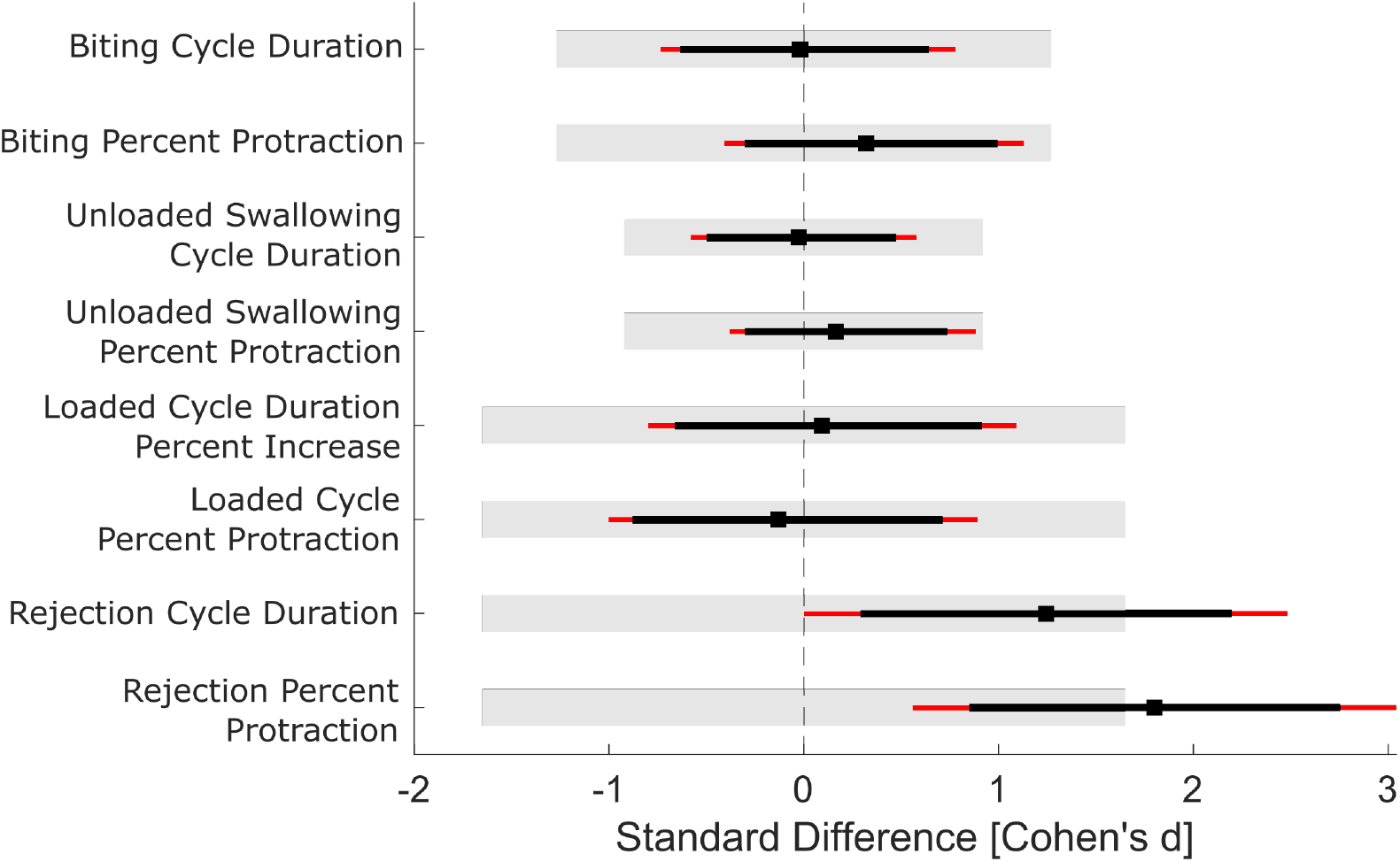
Statistical comparison between model predictions and animal data. Square marks show the normalized mean difference; black lines show the 90% Confidence Interval; and red lines show the 95% confidence interval. The gray-shaded region indicates the equivalence bound for the given metric. Positive standard differences indicate that the mean animal value is greater than the model value. Equivalence is supported if the 90% confidence interval is fully contained within the equivalence bound, which was true for all but the two rejection metrics. If the 95% confidence interval intersects the 0 difference line (dashed gray line), then the t-test fails to reject the null hypothesis.

### Comparison with previous biomechanical model

We performed a quantitative comparison with the previous state-of-the-art biomechanical model [21] to investigate if the additional computational complexity, and biological realism, of our new model provided substantial improvements (Fig. 10). To do this, we calculated both the cross-correlation and RMSE for the present and previous models relative to the animal’s internal kinematics. However, because the previous model had no geometry, this comparison could not be conducted in real-world units. Instead, the odontophore translation was normalized to the range of motion of each behavior, and these normalized curves were used to compare the models. The previous model already captured biting behavior very well, with a cross-correlation of *R* = 0.997 and RMSE of 5.62%, and the new model performed similarly well (*R* = 0.993, RMSE = 9.63%). However, the new model improved the kinematics of both unloaded swallowing and rejection, with RMSE decreases of 5.63% and 16.03%, respectively, compared to the previous model. The new model also better predicts the ratio between biting and swallowing cycle durations, with a 0.2% error in the new model compared to a 14.7% error in the previous model. Finally, the new model underpredicts the rejection to biting timing ratio, with an error of -13.5% compared to the previous model’s 11.2% overestimation.

**Fig 10.**
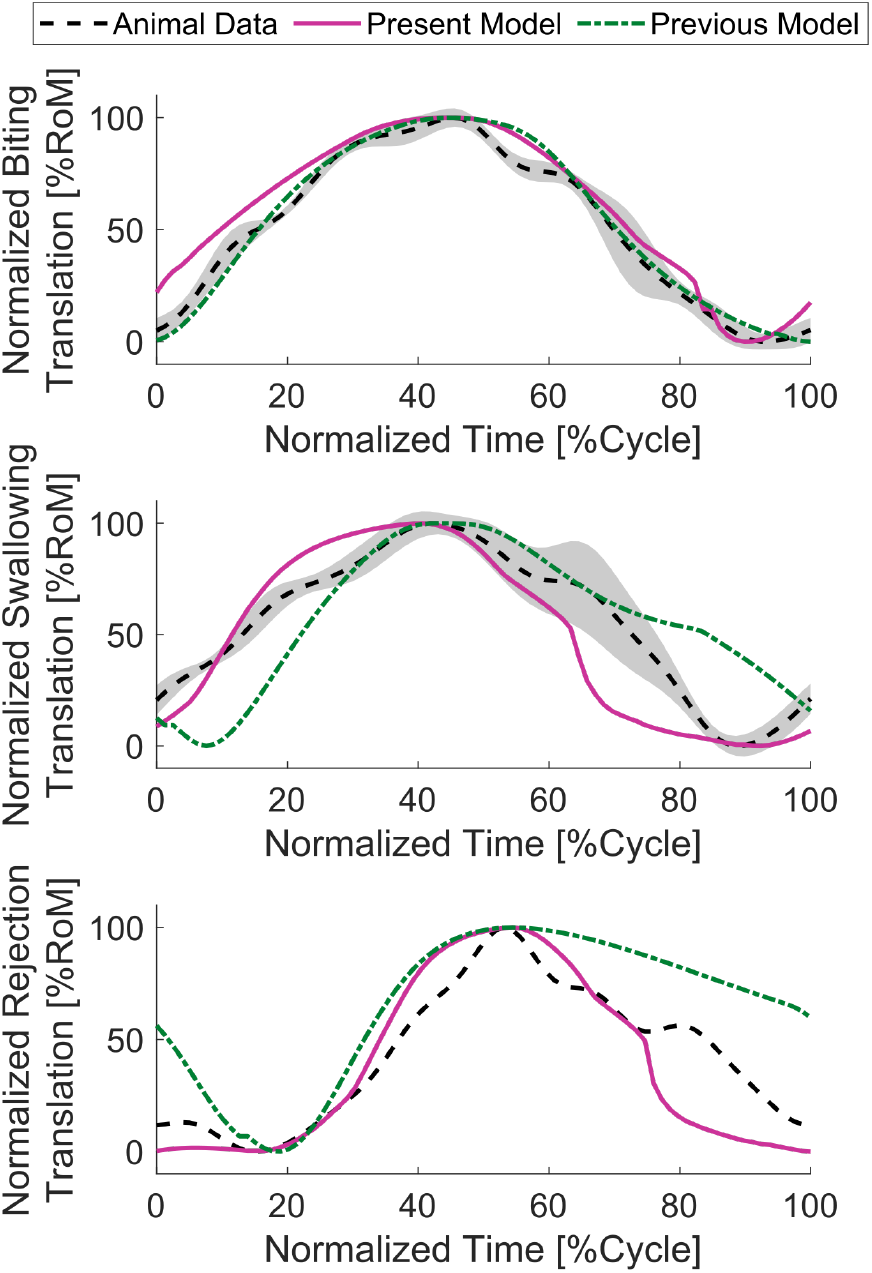
Comparison with previous biomechanical model. Animal data (biting and swallowing [32], rejection [23]) is presented as black dashed lines (standard deviation shown by the shaded region). Data from the new model is shown in the solid fuchsia line, and the previous model [21] is shown in the dot-dashed green line. Because the mechanics of the previous model only had nominal units, this comparison could not be conducted in real-world units. Instead, kinematic trajectories were normalized by the range of motion for each behavior. Both models capture the kinematics of biting very well (top), while the new model performs better in swallowing (middle) and particularly in rejection (bottom). Cross-correlation and RMSE values are reported in Table 4

### Time constants converge to *in vivo* measurements with no damping

During the tuning of the model, all parameters with units of time were normalized to the I2 time constant (i.e., in model units, *τ*_*I*2_ = 1). Then the value of the I2 time constant (in absolute units) was scaled such that the behavior durations of the model behaviors were in line with the times observed in animals. As all time-dependent values were uniformly scaled by this value, this acts only as a change of units and does not affect the dynamics of the system. Thus, based on animal behavior duration, we were able to make an independent prediction of the I2 time constant, in contrast to previous biomechanical models [21, 27] where the damping parameter was set at 1 and not tuned. However, during the tuning of this model, we discovered that to obtain the correct magnitude of translation in the correct amount of time (relative to animal data), the damping parameter had to be significantly lowered. For the main results presented here, that damping parameter was empirically set at 1/50 in model units. To examine how the model would behave in the no-damping limit, we conducted a sweep of damping parameters logarithmically spaced from 1.26 down to 0.0017 (Fig. 11). Below this value, the model experienced numerical instabilities due to the stiffness of the system and the specified tolerances of the solver. For each damping coefficient, biting behavior was simulated with an I2 time constant equal to 1, and the cycle time was calculated. Then, the required time scaling (and therefore the predicted value of *τ*_*I*2_) was calculated as the mean animal bite time divided by the measured cycle time. As the value of the damping coefficient approached 0, the predicted value of *τ*_*I*2_ approached a value of 1.1 s. From our fitting of the double-first-order filter model to the I2 activation function found in [35], we calculated a double-first-order filter time constant of 1.16 s for the animal data. This is in very good agreement with the value converged to by the model. This supports the idea that this model could be used to predict biological parameters by fitting to gross kinematic measures. Additionally, it provides additional evidence that damping does not play a significant role in shaping behaviors in this system. Because of this, damping may be removed in future iterations of this model, reducing the “dynamics” to a 0^th^ order system, as has been proposed previously [52].

**Table 4.**
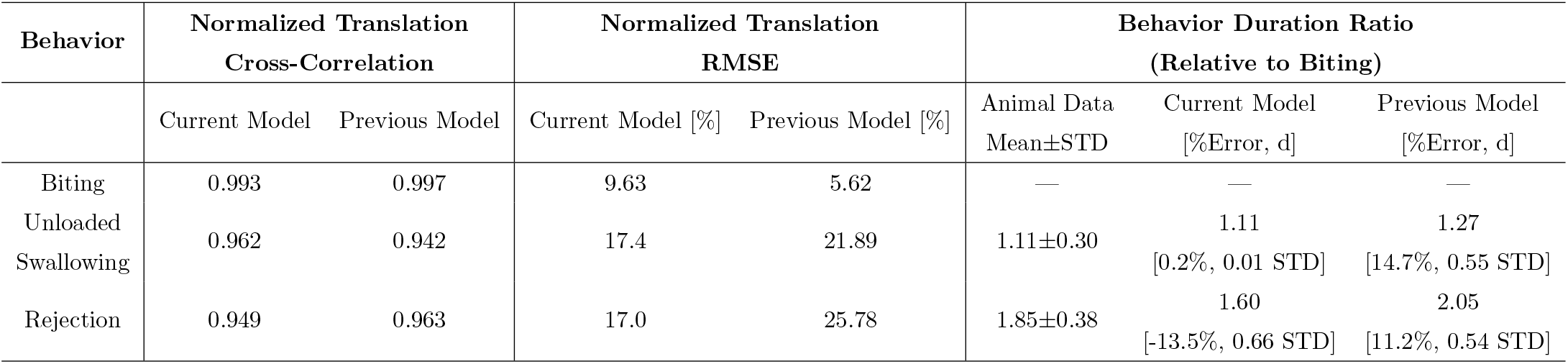
Quantitative comparison to the previous biomechanical model presented in Webster-Wood *et al*. 2020 [21].

**Fig 11.**
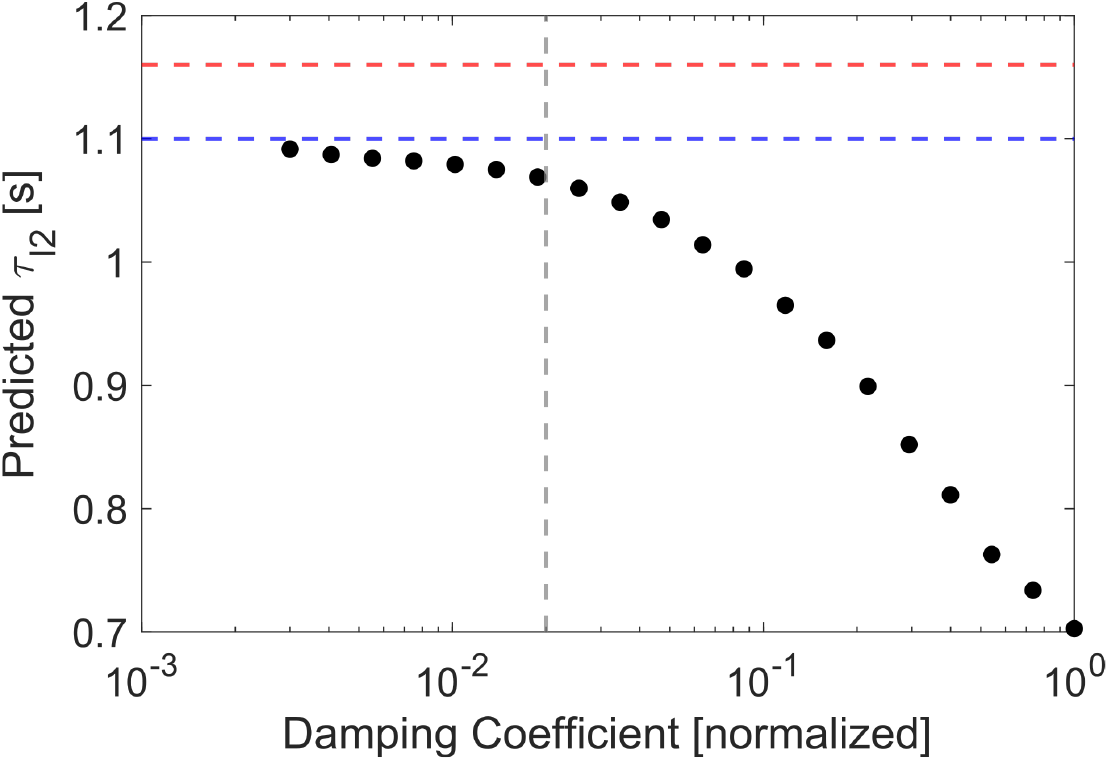
Convergence of time scaling with decreasing damping parameter. As the magnitude of the damping parameter, plotted in normalized model units, decreases towards 0 (no damping), the predicted value for the I2 time constant converges to a value of ∼ 1.1 s. Black circles show the results of individual simulations. The red dashed line shows the measured value of the I2 activation time constant of 1.16 s [35] (See Supporting Information 2.1). The blue line shows the converged value of the predicted I2 time constant. Finally, the vertical gray line shows the level of damping that was chosen for the final tuning.

## Discussion

The model presented here was largely successful in both qualitatively and quantitatively matching animal behavior. The three primary feeding behaviors demonstrated by the animal – biting, swallowing, and rejection – could be reproduced by switching the sensory cues provided to the model (Fig. 5), and the kinematics of these behaviors generally agreed well with internal kinematics measured in the animal (Fig. 7(c)). The model can also automatically switch between behaviors when these sensory cues are changed (Fig. 6). Compared to previous biomechanical models that were tuned solely to achieve multifunctionality, here, the model achieved both multifunctionality and quantitative agreement with animal data. This quantitative agreement to kinematics data was similar to or better than previous biomechanical models when comparing normalized data (Fig. 10, [21]). Because this new model incorporated explicit geometry, much closer to the real kinematics of the buccal mass, comparisons to animal data could also be made in real-world units (Figs. 7(c) and8, Table 3). The scalar metrics observed from the model were statistically equivalent to the measured animal data for both biting and swallowing behaviors (Fig. 9), though the model underpredicted the timescale of rejection behaviors.

Through tuning this model to kinematics and timing data, an independent estimate of the I2 time constant could be made, with this estimate agreeing with animal data within 5.5% (Fig. 11, [35]). The agreement with animal data was achieved using both parameter estimation to existing animal datasets and *ad hoc* hand-tuning. Even better performance could be achieved using numerical optimization techniques [62, 63].

## Limitations

### Choice of neural and muscle model

In our model, the neurons are modeled as binary switches instead of as continuously varying dynamical systems. This simplified model was successful in producing steady-state behaviors as demonstrated here and previously [21, 24, 27], but it is limited in its ability to investigate the effects of non-bursting neural signals in control [46], intermediate behaviors not coded in the cerebral neuron layer, such as rasping [26], and dynamical processes like neuromodulation [25, 64, 65]. It also cannot replicate the variability in the neural signals, which leads to cycle-to-cycle variability observed in animals [28, 30, 57]. Finally, the Boolean model’s ability to only capture “fully on” or “fully off” neural activity may be contributing to some of the mechanical disagreement in the model. In multiple behaviors, the model-predicted protraction is occurring too quickly. While this is likely predominantly related to the failings of the hinge and the lack of a passive I3 force (see “Muscles are more than active line elements”), this could also be related to an overestimation of the I2 force. The I2 motor neurons fire at between 12-17 Hz during *in vivo* swallowing behaviors [26], but the maximal force in I2 is not achieved until closer to 25 Hz stimulation [35]. Therefore, we would not expect the I2 to generate its maximal force during swallowing behaviors. Because the maximal force of I2 in the model was set using the maximal force measured in Yu *et al*. 1999 [35], a Boolean input of 1 would cause the I2 muscle to approach its maximal force, rather than an intermediate force associated with a stimulation frequency of ∼ 15 Hz. Utilizing a neural model with spiking neurons or a continuous rate-coded output may help alleviate these issues.

In addition to the simplified neuron models, our muscle models lack key features of biological muscles, including force-length and force-velocity relationships [66]. Such properties have been characterized in a limited number of *Aplysia* muscles [35, 37], but previous models indicate that they have important behavioral implications [22, 23].

Given the observed convergence to 0^*th*^ order dynamics in the biomechanics (Fig. 11), it may be possible to neglect the force-velocity properties of these muscles, modeling only their force-length properties and activation dynamics.

### Muscles are more than active line elements

While the model produces good quantitative matches to the animal data overall, there are still areas where improvement is needed to match the full cycle kinematics and neuromechanics better. First, it was observed that the rate of protraction was too fast: the slope of the odontophore translation curve is too high during protraction for swallowing and rejection behaviors. Although the range of motion in the behaviors was in good agreement with the animal data, the model produced this translation in a shorter amount of time than the animal, as shown in the higher Percent Protraction metric for the animals. Next, in both biting and rejection behaviors, the hinge was ineffective at producing retraction, and the model had to rely on the I3 to generate active force to compensate. While this produces kinematics that agree with the animal data, we know that in the animal, the I3 is not the sole retractor. In biting behaviors, while there is some activity in the I3 motor neurons, the frequency of firing is insufficient to generate large forces [46]. The hinge produces forces during biting protraction that are sufficient to retract the odontophore [36]. In rejections, the firing of B3/B6/B9 is delayed and the hinge muscle initiates retraction [58]. Finally, in both behaviors for which we have animal data, the model grossly underestimates the rotational range of motion of the odontophore. This is because, while the odontophore is free to rotate inside the I3 lumen, there are no force-generating elements that have sufficient mechanical advantage to generate this rotation until the odontophore has exited the lumen and can effectively “fall” backward. These discrepancies all provide evidence against assumptions A2 and A3 and point to two major mechanical shortcomings in the model. First, in agreement with assumption A3, the model I3 currently generates no passive forces other than the inequality constraints that act to resist protraction. In reality, the odontophore has to push open and stretch the I3 muscle to protract. These passive forces would help to slow protraction and assist in retraction without requiring the I3 to generate force actively. The second shortcoming is the general inability of the hinge to produce significant force. Currently, in agreement with assumption A2, the hinge is modeled as a single line-element muscle, but because the ventral I1 can shorten as the odontophore enters the I3 lumen, the hinge stays very near its rest length throughout feeding. This means it can generate very little force and exhibits minimal mechanical advantage in the system. Conversely, in the animal, the hinge is not only stretched but is also bent as the odontophore rotates into the I3 lumen, acting akin to a beam structure. This bending generates a reaction torque, both working to resist protraction and to generate additional rotation while the odontophore is within the I3 lumen. Additionally, as the hinge contracts, the bending stiffness of the hinge will increase, subsequently increasing the restoring torque on the odontophore and further assisting retraction. These simplifying assumptions were put forward following a demand-driven complexity analysis of the system in attempts to improve the model’s anatomical accuracy while minimizing the increase in computational complexity.

However, the results suggest that this simplification is inadequate to fully model the system, and in future versions of the model, the passive stiffness of the I3 lumen and the bending stiffness of the hinge should be incorporated.

### Need for a shape-changing odontophore

Finally, one significant simplification in this model is the assumption that the odontophore has a constant shape. This simplification (or further reductions of the odontophore to a point mass) has been utilized in multiple previous models of the system [21, 22, 52]. During the demand-driven complexity analysis, this assumption was considered sufficient, and therefore, it was not worth the additional computational cost to incorporate a deformable grasper. However, in the animal, the odontophore changes shape throughout the feeding cycle across all three major axes [32].

It has been hypothesized that this shape change influences the mechanical advantage of the different muscules in the buccal mass at different stages of feeding behavior [22, 23]. These hypotheses cannot be addressed in this model because of the fixed shape of the odontophore and lack of a length-tension model for the muscles [35]. The lack of a shape-changing odontophore also impacts the model observables and how they are compared to the animal data. This happens because there are additional internal degrees of freedom in the odontophore that are not captured in the model.

Translation in the animal is measured as the distance from the jaw line to the anterior edge of the odontophore [32]. In the model, this becomes simply an affine function of the center of mass position and head position. However, in the animal, as the odontophore deforms, the anterior edge may move, even if the center of mass does not. These deformation-based “translations” cannot be captured by the model. Additionally, radular stalk rotation in the animal is measured through a combined angle from the jaw line to I6 and from I6 to the radular stalk. As with translation, this becomes an affine function of the center of mass angle in the model. However, in the animal, as the odontophore changes shape, both the anterior line of the I6 and the radular stalk can move within the odontophore. These changes cannot be fully captured computationally without similar internal degrees of freedom or ways to calculate appropriate analogs.

For a more complete treatment of the system, a shape-changing odontophore should be incorporated and is likely to significantly improve the match between animal data and the model predictions, which could justify the additional computational cost.

## Acknowledgements

This work was supported by NSF DBI2015317 as part of the NSF/CIHR/DFG/FRQ/UKRI-MRC Next Generation Networks for Neuroscience Program. MJB and ASL were supported by the NSF Graduate Research Fellowship Program under Grant No. DGE2140739. Any opinions, findings, conclusions, or recommendations expressed in this material are those of the authors and do not necessarily reflect the views of the National Science Foundation. MJB was partially supported by faculty startup funding from the Carnegie Mellon University Mechanical Engineering Department.

## Author Contributions

**Conceptualization:** MJB, ASL, RS, BK, SMR, JPG, JMM, GPS, HJC, VWW

**Data curation:** MJB, JPG

**Formal analysis:** MJB

**Funding acquisition:** VWW, HJC, GPS

**Methodology:** MJB, ASL, RS, VWW

**Project administration:** MJB, VWW, HJC

**Resources:** VWW

**Software:** MJB

**Supervision:** VWW, HJC, GPS

**Visualization:** MJB

**Writing – original draft:** MJB, VWW

**Writing – review and editing:** MJB, ASL, RS, BK, SMR, JPG, JMM, GPS, HJC, VWW

## Supporting Information

**S1 Text. Computational methods for model observables, parameter estimation, and statistical analysis**. In this supporting information, we provide additional details related to the calculation of model observables, the process of model parameter estimation, and the statistical analysis performed in this paper. S1 Text contains five figures and three tables. Fig S1 shows the results of estimating the time constant of the I2 muscle based on the model presented in Yu *et al*. 1999 [35]. Fig S2 presents model fits for the time constants of the I4, hinge, and anterior I3 muscles using data reported in [49], [36], and [51] respectively. Fig S3 shows the resulting bootstrapped distributions for the animal cycle durations for biting, unloaded swallowing, and loaded swallowing. Fig S4 shows the resulting bootstrapped distributions for the Percent Protraction metric for the same behaviors. Finally, Fig S5 reports the results of the computational power analysis performed on the Equivalence Test for the different available sample sizes. The tables in S1 Text provide the parameters used in the model, with Tab 1 giving the mechanical model parameters, Tab 2 the neural threshold parameters, and Tab 3 additional (non-threshold) neural parameters.

