## Supporting Information 1 for "Incorporating buccal mass planar mechanics and anatomical features improves neuromechanical modeling of *Aplysia* feeding behavior"

#### 1 Model Observables Calculations

The following model observables were calculated and analyzed:

**Scalar Observables** :

1. **Cycle Time [s]**: the total cycle time was calculated as the time between sequential rising edges of the B31/B32 neurons
2. **Time in Protraction [s]**: the time spent in protraction is calculated as the duration of the B31/B32 motor pool firing, in agreement with the calculation performed in [2].
3. **Percent Protraction [%]**: the fraction of the cycle spent in protraction is the Time in Protraction divided by the Cycle Time.
4. **Translational Range of Motion [mm]**: the translational range of motion is calculated at the maximum value of odontophore translation (see below) minus the minimum value of odontophore translation.
5. **Rotational Range of Motion [deg]**: the rotational range of motion is similarly calculated as the maximum minus the minimum value of the odontophore angle (see below).

**Continuous Observables**

1. **Odontophore Translation [mm]**: following the measurements presented in [8, 9], the translation of the odontophore is not the center of mass motion, but is the distance from jaw line to the anterior edge of the odontophore, with the zero point corresponding to the odontophore crossing the plane of the jaws. This is calculated as:

$$\Delta x = R_{mm}\left[(x_g + R) - (L_{head} + x_h)\right] \tag{S1}$$

where $R_{mm}$ is the radius of the model odontophore in mm. See Parameter Estimation and Tuning.

2. **Odontophore Angle [deg]**: again following the metric reported in [8], the angle reported is not the centerline angle to the horizontal ($\theta_g$), but the angle of the radular stalk to the jaw line. Here, we estimate that the radular stalk axis aligns with the line from the center of mass $(x_g, y_g)$ to the food attachment point. Thus, the angle is reported as:

$$\theta_{obs} = \frac{180}{\pi}\left(\frac{\pi}{2} - (\theta_g - \theta_f)\right) \tag{S2}$$

3. **Force on Seaweed [nominal]**: the force exerted on the model buccal mass is the sum of the frictional force at the jaws and the grasper, such that:

$$F^{seaweed} = F^g + F^j \tag{S3}$$

4. **Length of Ingested Seaweed [mm]**: the amount of seaweed that the model ingested in a given swallow can be calculated by integrating the x-velocity of the food attachment point (relative to the jaw line) while the food is not slipping. Multiple conditions must be met for the food to be considered moving. First, the state of grasper slip must be 0 (See Frictional Forces), and the grasper must be sensing mechanical stimulus. Additionally, if the food attachment point is moving forward, then the jaws must be open (determined if $F^{I3_{anterior}} \leq 0.5F^{I3_{anterior}}_{max}$). If the jaws are closed, then the food does not move forward. This conditional integral of the velocity of the food attachment point was performed using an indicator function ($\delta_{ingest}$) that equals 1 when food was held by the grasper but not the jaws, and 0 otherwise:

$$L_{ingested}(t) = \int_0^t \delta_{ingest}(\tau)\dot{\tilde{x}}_f(\tau)d\tau \tag{S4}$$

where $\tilde{x}_f = x_f - (L_{head} - x_h)$ is the relative position of the food attachment point in the head, and the dot indicates the time derivative. The derivative $\dot{\tilde{x}}_f$ can be analytically calculated using the derivatives of $x_g$, $\theta_g$, and $x_h$. However, numerical stiffness associated with the discontinuous nature of the friction model caused a mean-zero jitter on these signals. This jitter caused issues for the direct use of the derivative signals because the condition regarding forward motion truncates the negative components of these jitters. This truncation biases the signal positive, leading to grossly overestimated values for the length of ingested seaweed. Therefore, $x_f$ is calculated from the output value of $x_g$, $\theta_g$, and $x_h$ and is smoothed with a 50-point moving average filter. The length of the filter was empirically chosen to be long enough to remove the jitter but short enough to maintain the full shape of the signal. The smoothed signal is then interpolated using a piecewise-polynomial spline from which analytical derivatives can be calculated. The derivative of this spline interpolation was then used in the integral above (Eqn. S4).

### 2 Parameter Estimation and Tuning

The model was tuned using a combination of data from the existing literature and hand tuning. Anatomical parameters for the model were estimated from the fixed buccal mass anatomy presented by [8] using ImageJ [10]. The radius of the odontophore was taken as half of the mean of the anteroposterior diameter and dorsoventral diameter. This was used as the reference length in the model, and all lengths were calculated in units of odontophore radii. This was also the conversion factor ($R_{mm}$) used to convert model units to absolute units in the Model Observables. The initial odontophore center of mass position was estimated as the intersection location between these diameters but was slightly adjusted to start the odontophore outside of the I3 lumen. The angle to the hinge $\theta_H$ was then calculated from the starting position using the pin-slot constraint to locate the hinge point. The angle to the food attachment point $\theta_f$ was found by drawing a line parallel to the radular stalk axis touching the anterior edge of the radular stalk and finding the point where this line intersects with the dorsal edge of the odontophore circle. The length of the I3 lumen was based on the projection of the dorsal and ventral I3 lines onto the horizontal. The I3 lumen length was used to set the initial values of $x_d$ and $x_v$. The height of the lumen $H_{lumen}$ was taken as the mean of the jaw line length and the lateral groove length. Using this rest position geometry, the rest lengths of all muscles were calculated as their length in this configuration.

The neuromechanical parameters of the model (muscle model parameters $\tau_j$, $T^j_{max}$, and $L^j_{max}$, passive spring and penalty stiffnesses, damping and frictional parameters, and neural model threshold parameters) were set through a combination of previously published literature and hand tuning. All parameter values can be found in Section 4. Values of $T^j_{max}$ and $\tau_j$ were normalized to the values for I2 such that, in model units, $T^{I2}_{max} = 1$ and $\tau_{I12} = 1$. Here, $\tau_{I2}$ is specifically the activation time constant of the I2 muscle. To determine the correct time I2 time constant for the double-first-order filter model, the time constant of this model was fit to simulations of the nonlinear activation function presented in [15] (See Section 2.1). This time constant was found to be 1.16 s. Given the number of elements in the system, there exists a large number of parameters to tune. Therefore, to reduce the parameter space dimensionality, the following simplifications were made.

1. It was assumed that all degrees of freedom experience the same level of damping ($C_q = \hat{C}$ for all linear degrees of freedom, and $C_{\theta_g} = \hat{C}/(2\pi)$).
2. Because the I4 and anterior I3 forces only impact the frictional forces and not any geometry parameters, it is not possible to meaningfully tune the static frictional parameters and the maximum tension parameters for the grasper and jaws independently. Therefore, the static frictional coefficient for both the grasper and jaws was set to 1, and the magnitude of the frictional force was fully determined by the maximum tension. The kinetic friction parameter was hand-tuned.
3. It was assumed that both I1 muscles and the bulk I3 had the same time constants, and this time constant was set at the mean of the activation and relaxation time constants found in [11].
4. It was also assumed that the dorsal and ventral I1 muscles generated the same maximum force and achieved the same maximum length. These parameters were hand tuned.
5. It was assumed that included extrinsic muscles have similar properties to each other, and thus E1, E2, and E6 were given the same time constant, maximum force, and maximum length relative to their rest length (i.e., the same maximum stretch).

Next, several parameters could be estimated from existing animal data. For I2, the maximum length relative to the rest length was estimated from [15] using the lengths at which the normalized passive force was 0 (at rest length) and 1 (at maximum length). Additionally, the maximum I2 force used to normalize all other force metrics was taken from [13]. The maximum force for I3 was also taken from [13] but was verified using the maximum force expected at physiological firing frequencies reported in [11]. The value from [13] is equal to the value predicted by [11] at a firing frequency of 12.6 Hz, well within the physiological range seen during behaviors [5]. Both the values of the maximum force and maximum length of the hinge were estimated from the [12] by observing the relative contributions of the active and passive forces when the hinge was protracted to the jaw line. Finally, the time constants for I4, the hinge, and the anterior I3 were estimated by fitting the double-first-order muscle model to stimulation response data reported in [7], [12], and [6], respectively.

The remaining parameters were first hand-tuned to produce multifunctionality and then refined to best match the available animal data, predominantly the internal kinematics. The conversion between model time units and absolute time (i.e., the I2 activation time constant) was then set to best match the cycle durations seen in animals.

#### 2.1 I2 Time Constant

To determine the appropriate time constant for the I2 muscle in the double-first-order filter model of muscle activation, this model was fit to simulations of the nonlinear activation function proposed in [15] (Fig. S1(a)). The nonlinear activation function is defined as

$$\frac{da}{dt} = \frac{1}{\tau}\left(u(t) - [\beta + (1-\beta)u(t)]\, a(t)\right) \tag{S5}$$

where $a(t)$ is the normalized level of activation in the muscle, $u(t)$ is the normalized neural input to the model, $\tau$ is the activation time constant, and $\beta$ is the ratio between the activation and relaxation time constant. In [15], $\tau = 2.45$ s and $\beta = 0.703$. To generate data from this model, simulations were run for 9 stimulation durations ranging from 3s to 27s. These data were the fit to the double-first-order filter model of the I2. As mentioned in the main manuscript, the I2 was given two time constants, one for activation and one for relaxation. Thus the governing equations for the model become,

$$\frac{d\tilde{T}^{I2}}{dt} = \frac{A^{I2}(t) - \tilde{T}^{I2}(t)}{\tau_{I2}^{k}} \text{ and} \tag{S6}$$

$$\frac{dA^{I2}}{dt} = \frac{N^{I2}(t) - A^{I2}(t)}{\tau_{I2}^{k}} \tag{S7}$$

where $\tilde{T}^{I2}$ is the normalized tension, $A^{I2}$ is the level of activation, $N^{I2}$ is the boolean neural input, and $\tau_{I2}^{k}$ for $k \in$ [a, r] are the activation and relaxation time constant respectively that is being fit. For a single squarewave stimulation at time $\hat{t}$ of duration $\Delta\hat{t}$, such that

$$N(t) = \begin{cases} 1 & t \in [\hat{t}, \hat{t} + \Delta\hat{t}] \\ 0 & \text{else} \end{cases} \tag{S8}$$

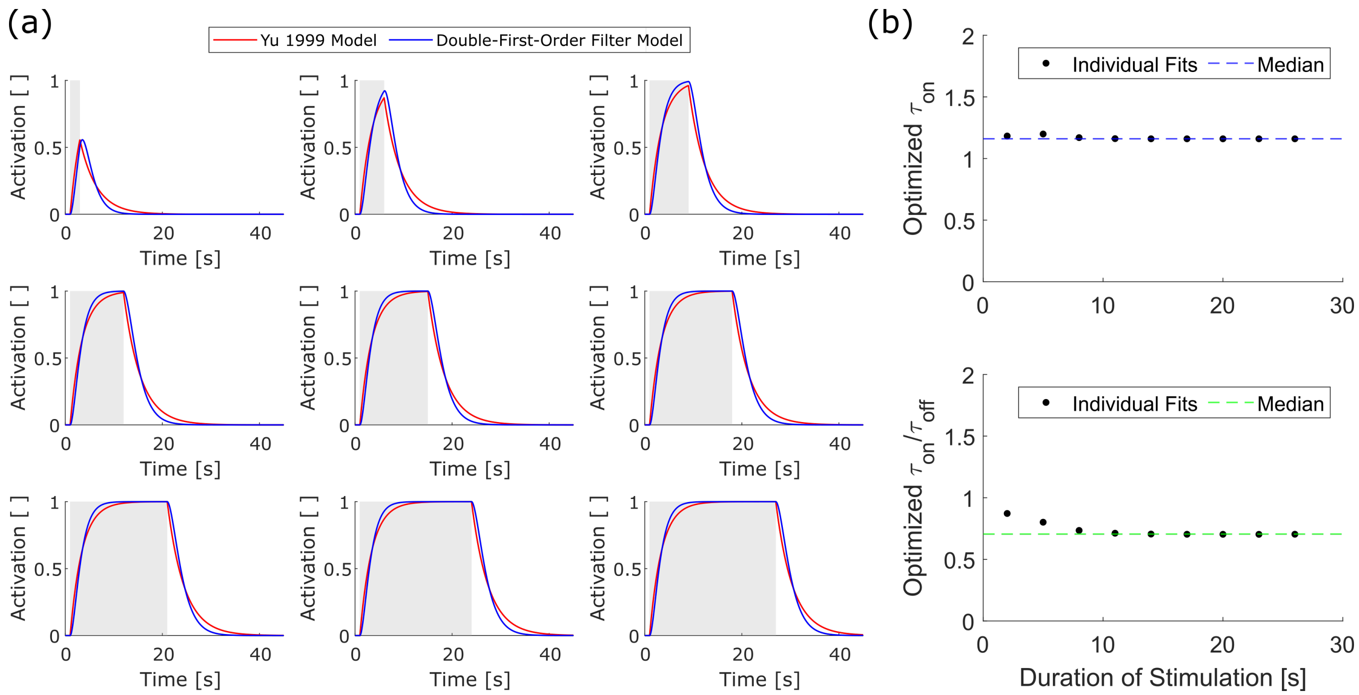


**Fig S1. Fitting of I2 time constant in the double-first-order filter to nonlinear activation function from** [15]. (a) Nonlinear activation function (red) and optimized double-first-order model (blue) for increase stimulation durations. The period stimulation is shown as the light gray box. (b) Parameter fits for the individual stimulation experiments, with the dashed line showing the median value that was utilized in the neuromechanical model.

the model tension response is solved as

$$T(t) = \begin{cases} 0 & t < \hat{t} \\ T_{rise}(t - \hat{t}) & t \in [\hat{t}, \Delta\hat{t}] \\ T_{fall}(t - (\hat{t} + \Delta\hat{t})) & t > \hat{t} + \Delta\hat{t} \end{cases} \tag{S9}$$

where

$$T_{rise}(t) = 1 - e^{\frac{-t}{\tau_j^{\mathrm{a}}}} \left(1 + \frac{t}{\tau_j^{\mathrm{a}}}\right) \tag{S10}$$

$$T_{fall}(t) = e^{\frac{-t}{\tau_j^{\mathrm{r}}}} \left(T_{peak} + A_{peak}\frac{t}{\tau_j^{\mathrm{r}}}\right) \tag{S11}$$

where $T_{peak} = T_{rise}(\Delta\hat{t})$ is the maximum achieved level of tension, and $A_{peak} = 1 - e^{\frac{-\Delta\hat{t}}{\tau_j^{\mathrm{a}}}}$ is the maximum achieved level of activation. $\tau_{I2}^{\mathrm{a}}$ and $\tau_{I2}^{\mathrm{r}}$ were fit by minimizing the squared error between $a(t)$ and $\tilde{T}(t)$ for each of the different stimulation durations. The final values of the time constants were then taken as the median time constant across all fits, which came to $\tau_{I2}^{\mathrm{a}} = 1.16$s and $\tau_{I2}^{\mathrm{a}}/\tau_{I2}^{\mathrm{r}} = 0.705$ (Fig. S1(b)-(c)).

#### 2.2 Time Constant Estimation

Time constants for I4, the I3 anterior, and the hinge were estimated from animal data by fitting the double-first-order filter muscle model to stimulation response data for that muscle (Fig. S2). The same closed-form solution to a squarewave stimulus presented in Eqs. S9-S11 was utilized in the parameter estimation. However, it was required that $\tau_j^{\mathrm{a}} = \tau_j^{\mathrm{r}}$ for these muscles. The value of both $\tau_j$ and $\hat{t}$ were fit to normalized data by minimizing the squared error using *fminsearch* (MathWorks). The value of $\Delta\hat{t}$ was measured from the animal data, but $\hat{t}$ was fit to account for unmodeled delays between stimulation and muscle response. The time constants for I4, the anterior I3, and hinge were fit to data from [7], [6], and [12], respectively. These time constants were then normalized by the activation time constant for I2, determined above, and these normalized time constants were used in the neuromechanical model.

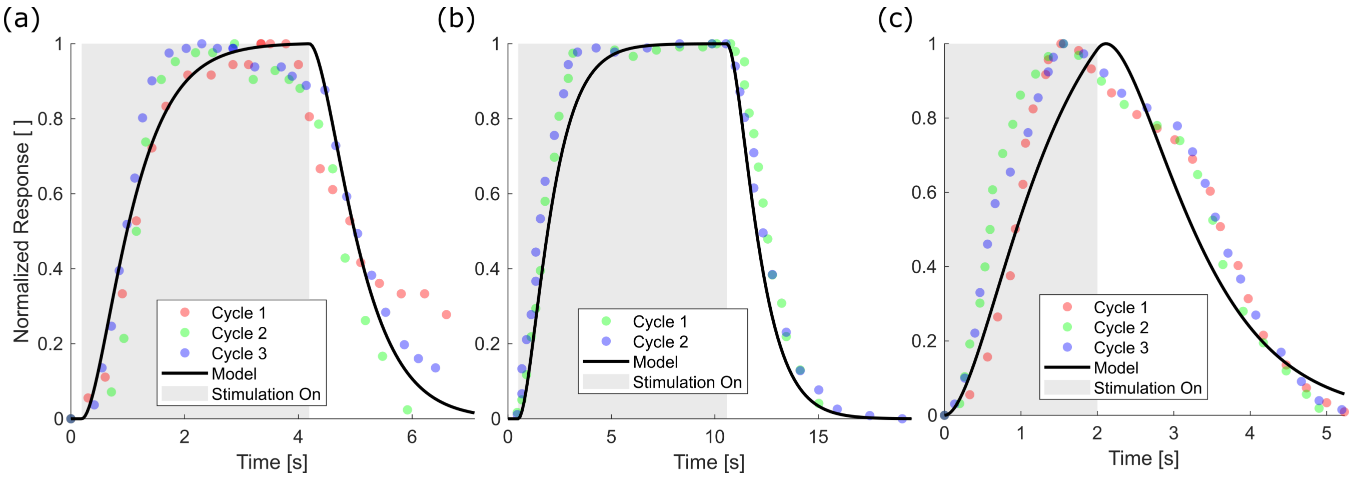


**Fig S2. Double-first-order muscle model fits to animal data.** Fits are shown for (a) I4 [7], (b) the hinge [12], and (c) the anterior I3 [6]. Each color marker shows an individual stimulation experiment, and the black line shows the double-first-order muscle response with optimized $\tau_j$ and $\hat{t}$. The light gray box shows when stimulation was applied to the muscle in question. The box begins at $\hat{t}$ and ends after the stimulation duration specified in the source study. All animal data were normalized by their peak value.

### 3 Animal Data and Statistical Methods

#### 3.1 Animal Data Bootstrapping

Scalar metrics were available for multiple individual animals for biting, unloaded swallowing, and loaded swallowing. For biting metrics, data were available for $N = 7$ animals from [2] (unpublished data); for unloaded swallowing metrics, data were available for $N = 12$ total animals between both [2] and [3]; and for loaded swallowing metrics, data were available for $N = 5$ animals. A bootstrapping approach was utilized to estimate the population mean of these metrics and to construct confidence intervals for hypothesis testing. This method also allowed for us to incorporate the uncertainties in the individual animal metrics into the population-level distribution.

For each animal, the datasets provided information to calculate a mean and standard error of the mean. This information was provided directly in the case of [3] and was calculated from individual cycle information in the case of [2]. The standard error captures the uncertainty in the mean value for a given animal due to the sample size and standard deviation in the metric. A sampling distribution for each animal's mean metric value can be constructed from this mean and standard error, with that sampling distribution being a normal distribution centered on the mean and with a standard deviation equal to the standard error. This gives a population of animals, each with their own sampling distribution for the mean metric of interest. To construct an estimate of the population distribution of mean values, a bootstrapped resampling of the animal population was conducted following the process laid out in Algorithm 1. For a behavior with a sample population of $M$ animals, $M \times 10,000$ resamples were taken.

**Algorithm 1** Animal Metric Bootstrapping

```
for i≤10,000M do
    Sample with replacement M animals from the sample population
    for each sampled animal j in the resampled population do
        Sample a mean value ~ N(μ_j, SE_j)
        μ̄_j ← sampled mean
    end for
    μ_booti ← (1/M) Σ_{j=1}^{M} μ̄_j
    σ_booti ← sqrt( (1/M) Σ_{j=1}^{M} (μ̄_j − <μ_booti>)^2 )
end for
```

$$\mu_{booti} \leftarrow \frac{1}{M}\sum_{j=1}^{M}\bar{\mu}_j$$

$$\sigma_{booti} \leftarrow \sqrt{\frac{1}{M}\sum_{j=1}^{M}\left(\bar{\mu}_j - <\mu_{booti}>\right)^2}$$

For metrics that are functions of the animal data instead of directly the animal data itself (i.e., protraction fraction, percent increase in cycle duration), at each step of the bootstrap, a mean is sampled from the sampling distribution for each input to the function. Then, the function is applied, and the result is stored. From the bootstrapped sampling distribution on the means ($\mu_{boot}$), we calculate the population average of the metric of interest as the mean

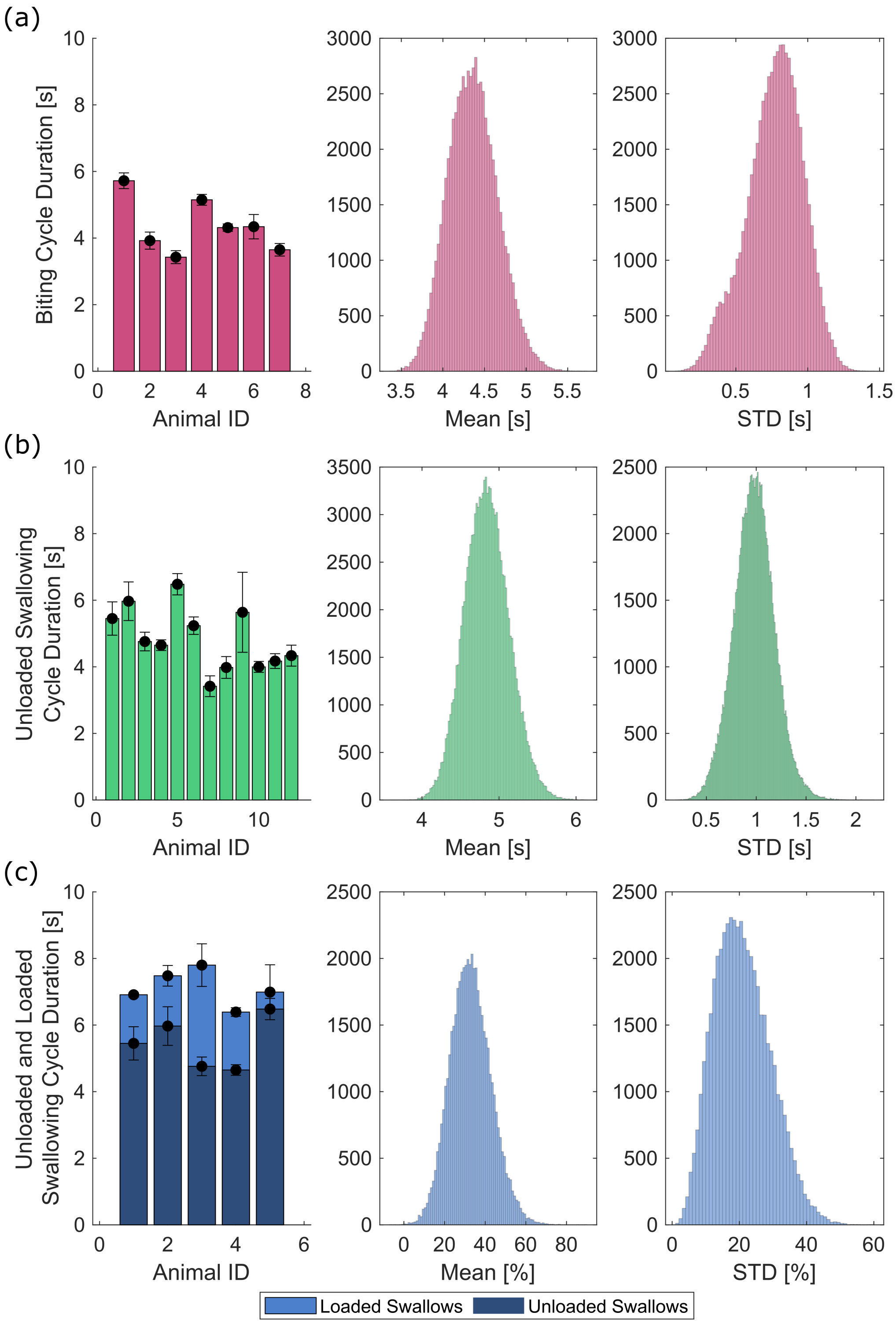


**Fig S3. Bootstrap distributions for (a) biting cycle duration, (b) unloaded swallowing cycle duration, and (c) percent increase in loaded swallowing cycle duration.** Bar charts show the mean value of an individual animal, and error bars show the standard error of the mean. These data were used as the parameters of the sampling distributions for each individual animal. The middle column shows the sampling distribution for the mean value of the metric, and the right column shows the sampling distribution for the standard deviation of the metric. From the mean sampling distribution, we calculate the population mean, standard error, and confidence intervals used in the equivalence tests. For (c), the dark blue bars show the cycle duration for unloaded swallowing, and the light blue bar shows the cycle duration of loaded swallows.

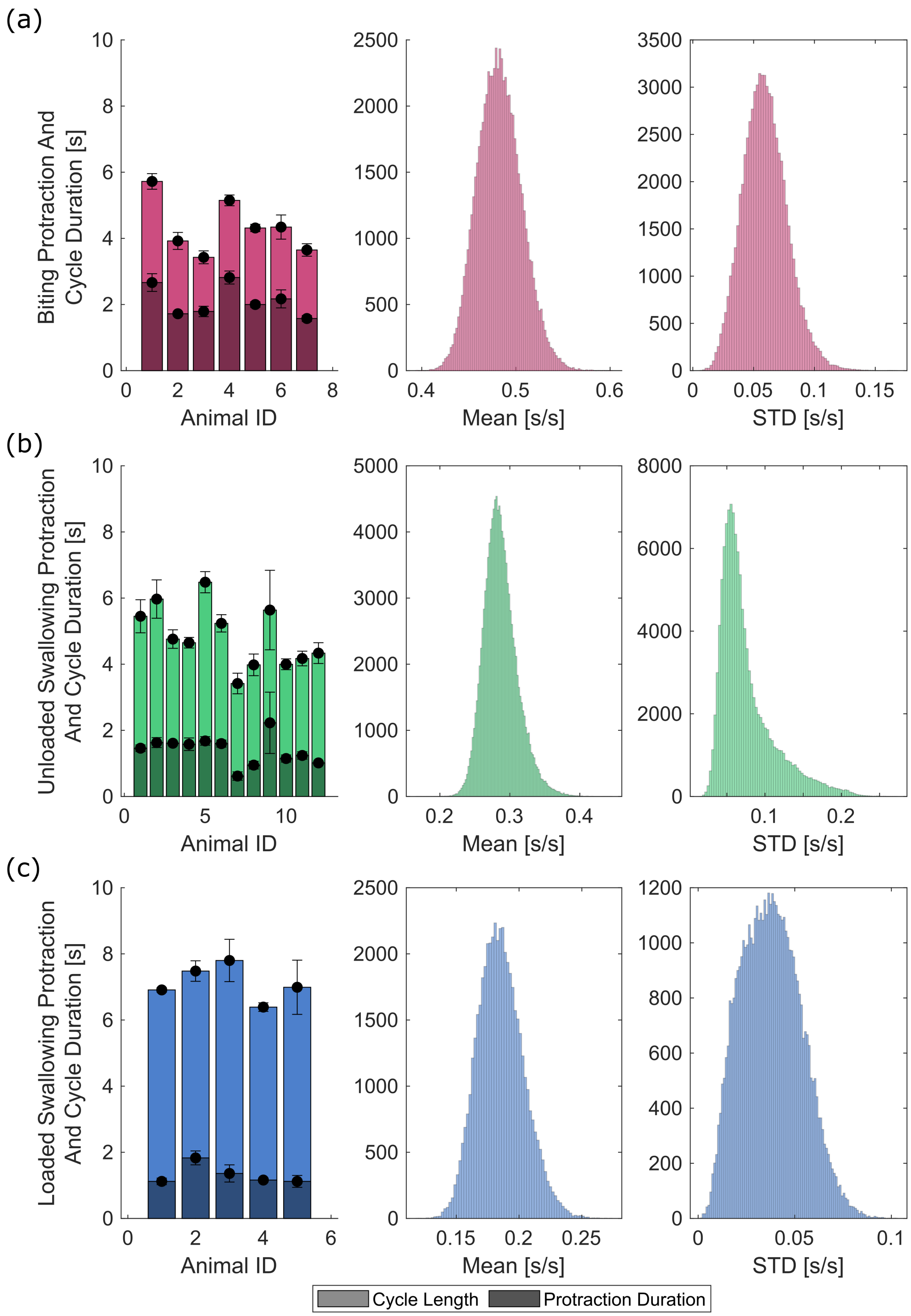


**Fig S4. Bootstrap distributions of protraction fraction for (a) biting, (b) unloaded swallowing, and (c) loaded swallowing.** In the left column, the light bar shows the mean cycle length and the dark bar shows the mean protraction duration for all three behaviors. Error bars show the standard error of the mean for each metric. The middle column shows the sampling distributions for the mean, and the right column shows the sampling distributions for the standard deviation. The mean sampling distributions were used to calculate the population mean, standard error, and confidence intervals.

value. We can also calculate the 90% and 95% confidence intervals on the metric from the quantiles of the sampling distribution. The population standard deviation was calculated as the mean of the standard deviation sampling distribution.

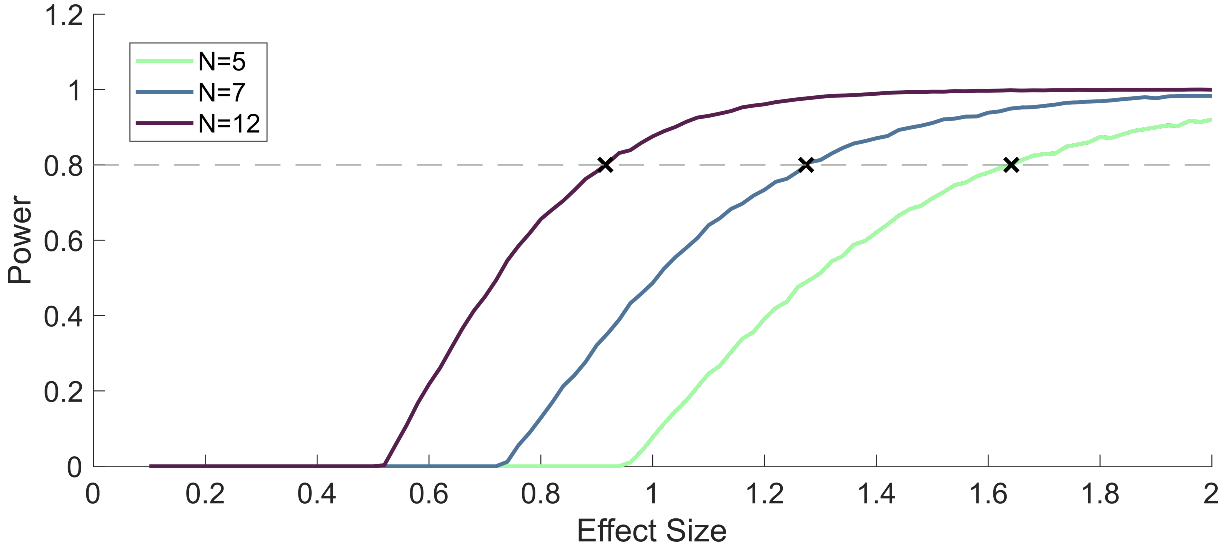


**Fig S5. Computational power analysis for the Equivalence Test at fixed sample sizes and varying equivalence intervals under the TOST procedure.** Power was calculated from 10,000 simulated experiments with a true difference of 0 for each effect size and sample size. The dashed gray line shows the target power of $1-\beta = 0.8$, and the black 'x' shows the computed intersection with the power curves for the different sample sizes.

All bootstrapping was conducted using custom MATLAB code and is available in the same repository as the main model code. For reproducibility, the pseudorandom number generator seed was set to 0 at the beginning of all random samplings [1]. The results of these parameter bootstraps are included below.

#### 3.2 Equivalence Test Power Analysis

To determine an appropriate equivalence interval based on the available animal data, a power analysis was conducted on the equivalence test. For a given sample size of animals, the equivalence interval was set at the effect size that could be detected at $1-\beta = 0.8$ for a true difference of 0 [4]. This power analysis was performed computationally using the following procedure [1]. As with the bootstrapped resamplings performed in Section 3.1), the pseudorandom number generator was given a seed value of 0 for reproducibility [1]. For a given sample size N and effect size d, 10,000 simulated experiments were run, For each experiment, N values were drawn from the animal population sampling distribution. Here that sampling distribution was set to a normal distribution with a mean and standard deviation of 1. However, because the Equivalence Test deals with standard differences and effect sizes, the absolute value of these parameters does not affect the power. The true difference was set to zero by setting the model mean value also to 1. Then an equivalence test was conducted on the sampled animal data and fixed model mean using the Two One Sample T test (TOST) protocol with an equivalence bound of ±d [4]. If the test succeeded (equivalence was observed at the $\alpha = 0.05$), an indicator variable for that experiment was set to 1; otherwise, the indicator variable was set to 0. This resulted in 10,000 indicator variables for a given sample size and effect size. The power for that sample and effect size is then calculated as the mean of the indicator function. Finally, to find the effect size that would yield a power $1-\beta = 0.8$ for a given sample size, the calculated power values were linearly interpolated and the point that minimized the squared distance between the interpolated values and 0.8 was calculated using *fminsearch* (MathWorks). These effect sizes for loaded swallowing and rejection ($N = 5$), biting ($N = 7$), and unloaded swallowing ($N = 12$) were found to be $d = 1.65$, $d = 1.27$, and $d = 0.92$, respectively.

#### 3.3 Exclusion of Loaded Swallowing Raw Cycle Times

When comparing scalar metrics between the model and animal data, it was decided not to compare the raw loaded swallow cycle times, but instead to compare the ratio of loaded to unloaded swallow cycle times. This is because the animals for whom we had these data skewed to the high end of the unloaded swallowing cycle time distribution, and thus may not be representative of the full population. We were able to observe this skewing because we have both unloaded and loaded swallowing data from the same animals from Gill *et al.* 2020 [3]. The animals from Gill *et al.* 2020 are labeled as Animal IDs 1-5 in Fig. S3 (b). The animals from this study had mean cycles times larger than all but two of the seven animals from Cullins *et al.* 2015 [2]. While this could be due to random statistical sampling (this cannot easily be verified with a larger independent sample of animal data), this is most likely due to the experimental conditions of the study. In Gill *et al.* 2020, the authors investigated the neural adaptation of *Aplysia* to a changing mechanical load during feeding. However, in order to perform these measurements, hook electrodes were surgically implanted. Therefore, all animals in this study underwent anesthesia and surgery prior to the behavioral studies. While all animals reported in the study successfully recovered and behaved similarly to previous studies, they may have been performing slightly less optimally than animals who had not undergone surgery, resulting in a slower cycling time. In our current model, we do not have a way to capture the effects of surgery on behavior, and therefore this effect cannot be controlled for. However, it was assumed that all behaviors were affected similarly by the surgery, such that the loaded and unloaded swallow times would have the same ratio as in the population as a whole. Therefore, we compared this metric to the data from our computational experiments.

### 4 Parameter Tables

The following parameters were utilized to perform all simulations presented in the main text. They are divided into parameters associated with the muscle and mechanical models (Tab. 1), those controlling neural feedback circuits (Tab. 2), and those controlling time delays in the neural model (Tab. 3).

**Table 1. Mechanical Model Parameters**. Here, the mechanical parameters of the model are summarized. All force values are reported in normalized model units (MU), and lengths are reported in units of grasper radii (R). In the definition of the max length, $L^j_{max}$, $L^j_0$ is the rest length of the same muscle, calculated as the length of the muscle in the rest configuration. The Source column indicates the source of the parameter (or data used to fit the parameter). HT indicates that the parameter was hand-tuned. The Passive Structure, Frictional, and Damping Parameters were all hand-tuned.

| **Muscle Parameters** | | | | | | |
|---|---|---|---|---|---|---|
| Muscle Name | $L_{max}$ [Grasper Radii] | Source | $T_{max}$ [Normalized] | Source | $\tau$ [s] | Source |
| I1 (dorsal and ventral) | $1.45L_0^{I1j}$ | HT | 3.2 | HT | 0.9998 | [11] |
| I2 | $1.6L_0^{I2}$ | [15] | 1 | — | 1.065 (on)<br>1.4182 (off) | [15] |
| I3 | — | — | 3 | [13, 11] | 0.9998 | [11] |
| I3 (anterior) | — | — | 0.1 | HT | 0.6256 | [6] |
| I4 | — | — | 0.9 | HT | 0.4362 | [7] |
| Hinge | 1.919 | HT | 0.292 | [12] | 0.780 | [12] |
| E1/E2/E6 | $1.5L_0^{Ej}$ | HT | 0.2 | HT | 0.606 | HT |

| **Passive Structure Parameters** | |
|---|---|
| Structure Name | Stiffness |
| Head Spring | 2 [MU/R] |
| Esophagus | 0.01 [MU/R] |
| Translation Constraint ($k_W$) | 10 [ MU/R] |
| Rotational Constraint ($k_{pen}$) | 0.5 [MU R / rad] |

| **Frictional Parameters** | |
|---|---|
| Friction Location | $\mu_{kinetic}/\mu_{static}$ |
| Grasper | 0.95 |
| Jaws | 0.75 |

| **Damping Parameters** | |
|---|---|
| Associated Degree of Freedom | Damping Parameter |
| $x_g$, $y_g$, $x_h$, $x_v$, $x_d$ | 0.0213 [MU s / R] |
| $\theta_g$ | 0.0213/($2\pi$) [MU R s / rad] |

**Table 2. Neural Feedback Threshold Parameters**. Each specified Post-synaptic Neuron receives an excitatory input related to the Feedback Parameter. This excitatory input is equal to the value of the Boolean statement. These connections are then combined with the rest of the Boolean logic of the neural network. The thresholds associated with each neuron and each feedback parameter are specific to each behavior that is performed. The pressure feedback parameter ($\tilde{P}_g$) is normalized, so all thresholds are in $[0, 1]$. For the translation feedback parameter $\hat{x}$, a value of 1 corresponds to the front edge of the odontophore touching the jaw line. Because the odontophore protracts past the jawline in some behaviors, the threshold values for this parameter can be greater than 1. All threshold values were hand-tuned.

| Behavior | Post-synaptic Neuron | Feedback Parameter | Description | Boolean Statement |
|---|---|---|---|---|
| Biting | B31/B32 | $\hat{x}$ | Translation threshold (when B31/B32 is active) | (B31B32== 0) AND ($\hat{x} < 0.54$) |
| | | | Translation threshold (when B31/B32 is inactive) | (B31B32== 1) AND ($\hat{x} < 1.1$) |
| | | $\tilde{P}_g$ | Pressure threshold | $\tilde{P}_g < 0.15$ |
| | B64 | $\hat{x}$ | Translation threshold | $\hat{x} > 0.56$ |
| | B3/B6/B9 | $\tilde{P}_g$ | Pressure threshold | $\tilde{P}_g > 0.5$ |
| | B7 | $\hat{x}$ | Translation threshold | $\hat{x} > 0.80$ |
| | | $\tilde{P}_g$ | Pressure threshold | $\tilde{P}_g > 0.70$ |
| Swallowing | B31/B32 | $\hat{x}$ | Translation threshold (when B31/B32 is active) | (B31B32== 0) AND ($\hat{x} < 0.26$) |
| | | | Translation threshold (when B31/B32 is inactive) | (B31B32== 1) AND ($\hat{x} < 0.74$) |
| | | $\tilde{P}_g$ | Pressure threshold | $\tilde{P}_g < 0.25$ |
| | B64 | $\hat{x}$ | Translation threshold | $\hat{x} > 0.24$ |
| | B3/B6/B9 | $\tilde{P}_g$ | Pressure threshold | $\tilde{P}_g > 0.75$ |
| | B38 | $\hat{x}$ | Translation threshold | $\hat{x} < 0.5$ |
| Rejection | B31/B32 | $\hat{x}$ | Translation threshold (when B31/B32 is active) | (B31B32== 0) AND ($\hat{x} < 0.40$) |
| | | | Translation threshold (when B31/B32 is inactive) | (B31B32== 1) AND ($\hat{x} < 0.81$) |
| | | $\tilde{P}_g$ | Pressure threshold | $\tilde{P}_g > 0.09$ |
| | B64 | $\hat{x}$ | Translation threshold | $\hat{x} > 0.22$ |
| | B3/B6/B9 | $\tilde{P}_g$ | Pressure threshold | $\tilde{P}_g < 0.30$ |
| | B7 | $\hat{x}$ | Translation threshold | $\hat{x} > 0.95$ |
| | | $\tilde{P}_g$ | Pressure threshold | $\tilde{P}_g > 0.60$ |
| | B4/B5 | $\hat{x}$ | Translation threshold | $\hat{x} > 1.1$ |

**Table 3. Additional Neural Model Parameters**. The follow model parameters also impact the behavior of the neural model, but do not relate to proprioceptive feedback. Instead, they model to delays in the model related by refractory periods and slow excitatory connects. For full descriptions of the modeled mechanisms, see Webster-Wood 2020 [14]. The parameter values were hand-tuned and are different values than used in [14].

| Parameter Description | Parameter Value |
|---|---|
| Duration of CBI-3 refractory period | 6.4539 s |
| Duration of B8 excitation after B40/B30 cessation | 4.0470 s |

11. Ravesh Sukhnandan, Qianxue Chen, Jiayi Shen, Samantha Pao, Yu Huan, Gregory P. Sutton, Jeffrey P. Gill, Hillel J. Chiel, and Victoria A. Webster-Wood. Full Hill-type muscle model of the I1/I3 retractor muscle complex in *Aplysia californica*. *Biological Cybernetics*, jun 2024.

12. Gregory P. Sutton, J. B. Macknin, S. S. Gartman, G. P. Sunny, R. D. Beer, Patrick E. Crago, David M. Neustadter, and H. J. Chiel. Passive hinge forces in the feeding apparatus of *Aplysia* aid retraction during biting but not during swallowing. *Journal of Comparative Physiology A: Neuroethology, Sensory, Neural, and Behavioral Physiology*, 190(6):501–514, 2004.

13. Gregory P. Sutton, Elizabeth V. Mangan, David M. Neustadter, Randall D. Beer, Patrick E. Crago, and H. J. Chiel. Neural control exploits changing mechanical advantage and context dependence to generate different feeding responses in *Aplysia*. *Biological Cybernetics*, 91(5):333–345, 2004.

14. Victoria A Webster-wood, Jeffrey P. Gill, Peter J Thomas, and H. J. Chiel. Control for multifunctionality : Bioinspired control based on feeding in *Aplysia californica*. *Biological Cybernetics*, 114(6):557–588, 2020.

15. Sung Nien Yu, Patrick E. Crago, and H. J. Chiel. Biomechanical properties and a kinetic simulation model of the smooth muscle I2 in the buccal mass of *Aplysia*. *Biological Cybernetics*, 81(5-6):505–513, 1999.
